# *Ustilago maydis* infection reshapes the maize phyllosphere microbiome through antimicrobial effectors and host metabolic reprogramming

**DOI:** 10.64898/2026.03.27.714703

**Authors:** Zarah Sorger, Sarah Daher, Frowin Reichhardt, Bilal Ökmen, Nadine Töpfer, Gunther Doehlemann

**Author notes:** contributed equally (Zarah Sorger, Sarah Daher).

## Abstract

Plant-associated microbial communities play a critical role in plant health and disease resistance, but the mechanisms which reshape these communities during pathogen infection are poorly understood. In this study, we investigated how infection of maize by the smut fungus *Ustilago maydis* is functionally linked with the bacterial phyllosphere microbiome and explored the role of an antimicrobial effector GH25 in fungal infection. Using a combination of culture-dependent and culture-independent approaches, we compared the leaf microbiomes of infected and uninfected plants. We observed a significant increase in microbial abundance and pronounced shifts in community composition and identified distinct health-associated (HCom) and disease-associated (DCom) bacterial communities. To assess whether *U. maydis* directly manipulates the microbiome, we tested the antimicrobial activity of the antimicrobial effector GH25 against isolated strains. Notably, all HCom bacteria were sensitive to GH25 and co-inoculation of HCom bacteria with a *U. maydis* Δ*gh25* knockout mutant significantly reduced fungal virulence. In contrast, DCom exhibited minimal sensitivity to *U. maydis* and did not affect the virulence of *U. maydis* Δ*gh25*. Genome-scale metabolic community modelling coupled with functional profiling revealed infection-associated shifts in predicted metabolic potential, consistent with *U. maydis* induced leaf tumors being strong sink tissues. Together, this work shows that *U. maydis* infection reshapes the maize phyllosphere microbiome through a combination of effector-mediated antimicrobial activity and host metabolic reprogramming.

## Introduction

The importance of the plant microbiome for plant health has become undeniable^1–3^. Plants are colonized by a multitude of microorganisms, including fungi, bacteria, viruses and oomycetes, which can thrive on the plant surface and within the plant, both below- and above-ground. Plants can actively shape their microbiota through chemical signalling, metabolic selection, and immune regulation^4–6^. This involves the secretion of root exudates and secondary metabolites that selectively promote beneficial microbes while suppressing pathogens^7,8^. Coumarins and glucosinolates are plant-derived secondary metabolites that refine microbiome assembly by acting as selective chemical filters in the rhizosphere^9^. Beyond the rhizosphere, plants host diverse microbial communities in the aerial parts, collectively known as the phyllosphere. Within these communities, commensal microbes are associated with improvement of plant defence against pathogens, through upregulation of defence related genes^10^. Together, these processes allow plants to dynamically recruit and maintain a functional microbiome adapted to their physiological state and environmental conditions^11^. Recently, the plant microbiome has even been defined as an extension of the plant immune system, underpinning the high relevance of a stable microbiome for plant health^12^. Both root and shoot microbiomes contribute critically to plant defense and are interconnected through systemic signaling and immune-metabolic coordination^13–15^. Nevertheless, often pathogens overcome plant immunity, thus being increasingly recognized not merely as invaders but as active engineers of the host-associated microbiome.

Accumulating evidence indicates that pathogen infection can profoundly reshape host-associated microbiomes across plant species and tissue compartments. In rice, infection with the foliar pathogen *Magnaporthe oryzae* has been shown to lead to alterations of the microbiome residing in rhizosphere and root compartments^16^. Moreover, infection with bacterial wilt disease (BWD) led to alterations in community function and assembly of both rhizo- and endosphere in tomato plants^17^. In cotton, *Verticillium dahliae* infection altered rhizosphere composition and co-occurrence network structure i.e., pathogen infection reshapes rhizosphere taxa and interaction networks^18^. While most studies focus on modulation of compartments associated with the root, manipulation of phyllosphere microbiome has also been reported. Infection of maize with *Exserohilum turcicum* has been reported to lead to shifts in both microbiome diversity and function, showing a significant reduction in alpha diversity after infection^19^.

However, the mechanistic basis leading to the microbiome remodeling remains largely elusive. Recently, three major processes mediating this have been proposed: (i) effects of activation of the plant immune system like alterations of the local physiochemical niche, (ii) recruitment of beneficial microbes by the host, and (iii) direct manipulation of the host microbiome through the pathogen by antimicrobial effectors and niche competition^20^. Many studies largely focus on recruitment of beneficial microbes as a thriving force, also known as “cry-for-help” response, where modification of root exudation can lead to microbiome restructuring, ultimately promoting disease suppression^14,21^. Still, it is likely that a combination of all these factors, including the pathogen, plant species, and environmental context will influence the outcome. For example, pathogen infection can trigger plant signals that can alter the rhizosphere microbiota, both in the infected plant and in neighbouring plants via volatile signals, influencing the abundance and diversity of suppressive and non-suppressive microbes^22^. Moreover, pathogens influence microbe–microbe interactions, as shown in gnotobiotic *Arabidopsis thaliana* where *Pseudomonas syringae* challenge shifts the ecological balance within commensal communities, affecting plant protection^23^. Microbe-microbe interactions during pathogen infection are often mediated by pathogen-secreted effectors with antimicrobial activity. Many such fungal effectors have been shown to have ancient origins, indicating that this antagonism between microbes might predate pathogen-host interactions^24,25^.

Pathogen-induced tumors represent an extreme and underexplored display of these processes, as they combine immune modulation, host metabolic reprogramming, and direct microbial interference within a spatially confined and developmentally stable tissue niche. Here, we focus on the biotrophic smut pathogen *Ustilago maydis*, which infects *Zea mays* and induces the formation of large tumor-like organs that are characterized by profound developmental and metabolic reprogramming of host tissue^26–29^. Although *U. maydis* is a well-established model system for studying molecular mechanisms of plant–pathogen and biotrophic interactions^30,31^, the interaction between *U. maydis* and the microbial communities that occupy the same niche remains largely unexplored. A negative association with *Fusarium verticillioides* has been reported, where the reduction of virulence and tumor formation of *U. maydis* was observed in maize leaves^32^. Nevertheless, knowledge about microbiome manipulation through *U. maydis* is sparse. Possible mechanisms can, however, be postulated based on the available knowledge of closely related species that colonize the phyllosphere, as well as antimicrobial compounds secreted by *U. maydis*. Non-pathogenic yeasts closely related to smut fungi, such as *Pseudozyma* spp., commonly colonize plant surfaces and can act as biocontrol agents by producing antifungal metabolites and biosurfactants that suppress phytopathogens^33,34^. In addition, *U. maydis* produces ustilagic acid, whose biosynthesis is controlled by a dedicated gene cluster which exhibits antimicrobial activity, suggesting a role in microbial competition^35,36^. Importantly, *U. maydis* also actively interferes with other microbes through secreted effectors, as demonstrated for Ribo1, a conserved extracellular ribonuclease that enables smut fungi to compete with host-associated bacteria^37^. Together, these features make the maize–*U. maydis* interaction a uniquely tractable system to investigate how infection and pathogen-induced tumors may reshape host-associated microbiomes.

Here, we investigate how infection by *U. maydis* shapes the phyllosphere microbiome of *Zea mays* and report robust microbial restructuring during infection. Moreover, we examined the antimicrobial potential of a homolog of a glycoside hydrolase (GH25) described to be involved in interkingdom microbial antagonism in a closely related beneficial yeast^38^ and established GH25 as a crucial antimicrobial effector promoting *U. maydis* virulence. Together, our results demonstrate that infection by *U. maydis* reshapes the maize phyllosphere microbiome by combining effector-mediated antimicrobial activity with host metabolic reprogramming.

## Results

### *Ustilago maydis* infection reshapes the phyllosphere microbiome of *Zea mays*

Pathogen infection has been shown to shape the plant microbiome, leading to changes in microbial composition, diversity, and community function^39^. We specifically aimed to investigate the influence of the biotrophic smut fungus *U. maydis* on the microbiome composition of *Z. mays*. Inoculations of *Z. mays* with *U. maydis* were performed. Plants were grown in pots on the field with a mixture of field and commercial soil, to include microorganisms from both soil and air (**Fig. 1A**, **Fig. S1**). At 15 days post-inoculation (dpi), region of infection and corresponding region in non-infected samples were collected and further separated into different fractions: an epiphytic fraction, an endophytic fraction after the epiphytes were washed off, an additional endophytic fraction after surface sterilization, as well as a combined fraction (whole microbiome). For each fraction, colony forming units (CFU) assays were performed to assess differences in bacterial abundances after infection (**Fig. 1A,B**). Strikingly, infection led to a significant increase of colony forming units detected on LB medium after *U. maydis* infection compared to non-infected samples (**Fig. 1B, Fig. S2**), indicating a higher abundance of fast-growing bacteria. Furthermore, leaf imprints were taken from each fraction in isolation, revealing greater bacterial colony growth in the infected samples than in the non-infected ones (**Fig. S2**). To get a deeper understanding of the changes in microbiome composition, 16S Amplicon Metagenomic Sequencing was performed. For each fraction, a drastic shift in the community composition was observed on the order level (**Fig. 1C, Fig. S3**). While non-infected plants depicted a diverse microbiome and were not dominated by a specific order, infected plants showed a predominance of *Pseudomonadales* and *Enterobacteriales*, displaying a relative abundance of at least 0.75 for all infection fractions. A flower diagram of Amplicon Sequencing Variants (ASVs) sequenced in each fraction revealed a core community of 32 ASVs, being present in each fraction (**Fig. 1D, Dataset S1**). Moreover, higher counts for ASVs in non-infected fractions compared to infected fractions further underscores the higher diversity of the non-infected microbiome. Results presented indicate a strong reshaping due to infection and rather subtle differences between fractions. However, a taxonomic heatmap on the genus level comparing all fractions reveals that the epiphytic non-infected fraction has a higher abundance of e.g *Deinococcus* and *Cutibacterium* compared to other fractions (**Fig. 1E**). Alpha-diversity metrics differed between treatments and were accordant with inferred community assembly processes. Infected samples exhibited increased dominance (lower evenness), whereas non-infected leaves showed significantly higher Shannon diversity for each fraction (**Fig. 1F, Dataset S2**). Consistent with these patterns, iCAMP^40^ analysis indicated that communities in non-infected fractions exhibited a more balanced contribution of dispersal limitation (DL) and homogeneous selection (HoS), alongside a substantial role of drift (DR). In contrast, infected tissue was characterized by an increase in drift-driven assembly processes, with DR accounting for the vast majority of community turnover across most fractions, while the relative influence of DL and HoS was reduced (**Fig. 1G**).

**Figure 1:**
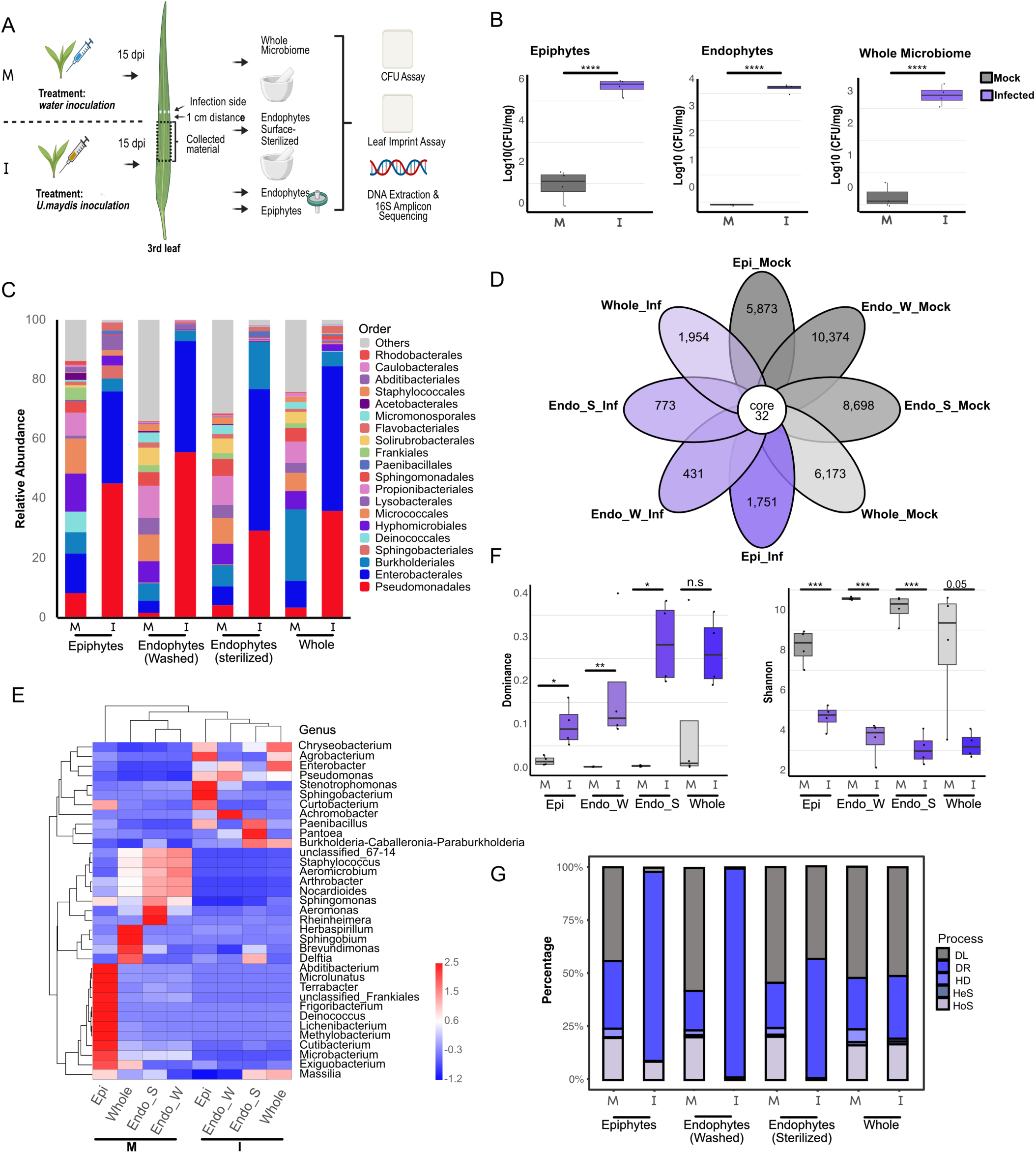
Infection with *Ustilago maydis* significantly alters microbiome composition. **A**) Overview of the workflow for the amplicon sequencing experiment: plants were inoculated with either water (mock) or *U. maydis*, and samples were collected at 15 dpi from the tumor tissue region. Four different fractions were isolated: epiphytic fraction, endophytic fraction after epiphyte wash (Endo_W), endophytic fraction after surface sterilization (Endo_S), as well as a whole microbiome fraction, where no separation occurred. A leaf imprint test and colony forming units (CFU) assay were performed before sample processing for amplicon sequencing. M: Mock treated, I: Infected **B**) Infection with *U. maydis* significantly increased the abundance of both epiphytes and endophytes in the phyllosphere. For endophytic samples, leaf material was taken, weighed and ground, whereas for the epiphytic samples, weight of total plant material was taken and part of the wash solution was used for the CFU assay. Dilution series was performed and plated on LB plates. Growth of strains was assessed by counting the amount of colony forming units and calculating the amount of CFU (Log10(CFU/ml)). **C)** Relative abundances of main groups revealed a shift in the microbiome after *U. maydis* infects the plant. The top 20 taxa of each group at the taxonomic rank “Order” were selected to form a distribution histogram of the relative abundance of these taxa. This allows the visualization of the taxa with a higher abundance and their proportion. **D)** Petal diagram of ASV distributions (for >5 samples/groups): Each petal represents a sample/group (different colors for different samples/groups). The “core” number in the center indicates ASVs shared by all samples/groups; numbers on petals indicate unique ASVs of the sample/group. **E)** Heatmap of taxonomic abundance: the color in each cell reflects the Z-score of the relative abundance of each row of species after standardization. As a result, colors in the heatmap can only be compared horizontally (composition of the same species across samples) rather than vertically (different species in the same sample). **F**) Representation of alpha diversity measurements, depicting both dominance and Shannon diversity. * Indicates statistical significance (Welch’s t-test) p < 0.05. **G**) This visualization presents the relative contributions of five microbial community assembly processes across experimental groups using a stacked bar plot format. The x-axis displays different treatment conditions or sample groupings, while the y-axis shows the percentage contribution of each assembly mechanism within respective groups. Color-coded segments represent: (1) homogeneous selection (HoS), (2) heterogeneous selection (HeS), (3) homogenizing dispersal (HD), (4) dispersal limitation (DL), and (5) ecological drift (DR), quantitatively demonstrating how dominant assembly patterns shift under varying environmental conditions. M: Mock treated, I: Infected. Figure 1A was created with Biorender.com

Taken together, *U. maydis* infection significantly reshapes the phyllosphere microbiome, both affecting epiphytic and endophytic communities. Next, we aimed to identify bacteria specific to fractions and conditions and functionally test how *U. maydis* infection might influence these bacteria directly or indirectly.

### Epiphytic community displays specific marker strains only present in uninfected plants

To functionally test whether *U. maydis* might actively shape the microbiome by the targeting of specific bacteria, we aimed to identify specific epiphytic marker strains for functional analysis. Combining culture-dependent and -independent approaches can result in the selection of specific potential target strains, which could be easily overlooked by simply relying on one approach. Bacterial isolation was conducted to create a culture collection (**Fig. 2A**). We isolated epiphytes and endophytes from infected and non-infected maize seedlings, as well as from adult plants. In total, 149 bacterial strains were isolated, of which the genus was determined using 16S sequencing (**Dataset S3**, **Fig. S4D**). With these strains at hand, we focused on epiphytic bacterial isolates. We hypothesized that these isolates form the first microbial barrier and are therefore the most likely direct competitors encountered by *U. maydis* prior to infection. In accordance with the CFU assay results (**Fig. 1B, Fig. S4A-B**), the abundance of strains that were culturable was increased after infection. We screened the isolated strains for epiphytic bacteria, which would only be observed in non-infected seedlings, as well as for isolates which were exclusively isolated from infected plant material (**Fig. 2B**). Eight strains were exclusive to non-infected plants, of which two were slow growing and thus removed from further analyses. The six remaining strains were subjected to full genome sequencing (see **Table S1** for summary of taxonomic identification using GTDB-Tk 2.7.1^41–47^ software and TYGS^48–55^ database), which identified them as *Priesta megaterium*, *Bacillus licheniformis*, *Paenibacillus amylolyticus*, *Staphylococcus epidermidis*, *Arthrobacter* sp., and *Peribacillus* sp. Epiphytic bacteria exclusively isolated from infected plant material were more diverse and generally more abundant, consisting of *Pseudomonas* sp.*, Enterobacter* sp.*, Curtobacterium* sp., *Paenibacillus* sp., and *Bacillus* sp., among others (**Dataset S3**). We restricted subsequent analysis to the following strains, which were also subjected to genome sequencing: *Enterobacter kobei, Brevibacterium sediminis, Microbacterium* sp.*, Erwinia aphidicola, Raoultella ornithinolytica,* and *Pseudomonas aestiva* (see **Table S1** for details). For further analysis, we grouped the six epiphytic strains exclusively isolated from non-infected maize seedlings into a “health community” (hereafter: “HCom”), as well as the six strains exclusively isolated from *U. maydis* infected plants into a “disease community” (hereafter: “DCom”).

**Figure 2:**
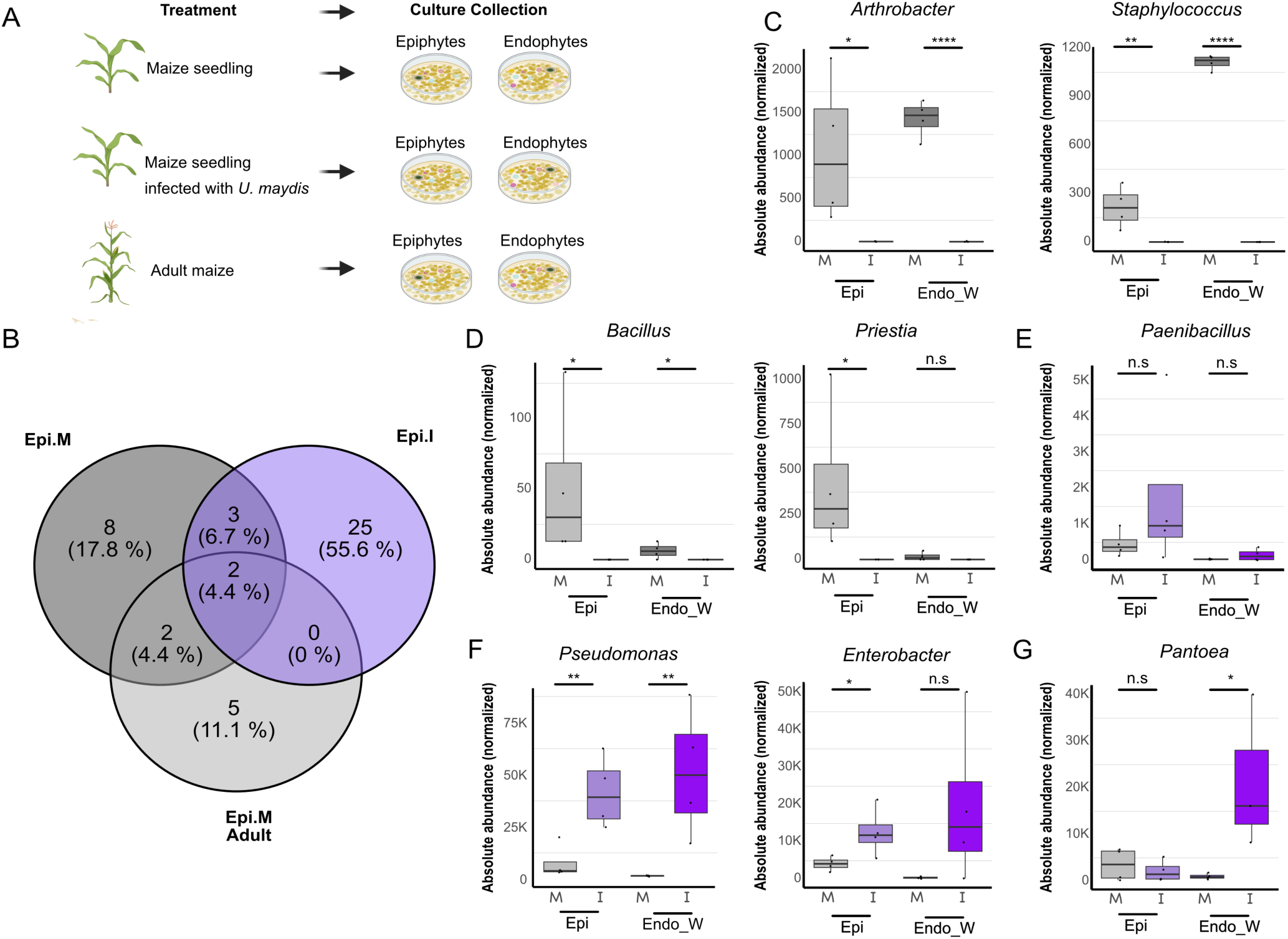
Comparison of culture-dependent and independent results for the selection of bacterial strains targeted by *Ustilago maydis.* **A**) A culture collection of epiphytic and endophytic bacteria was generated by using material from uninfected Golden Bantam (GB) maize seedlings (mock), maize seedlings infected with *U. maydis*, as well as adult maize plants (cultivar: Ky21). **B**) Isolated bacteria were screened for specificity to mock or infection fraction. “Epi.M” = epiphyte fraction from mock seedlings, “Epi.I” = epiphyte fraction from infected seedlings, “Epi.M.Adult” = epiphyte fraction from adult plants. **C**-**G**) Identified strains correlating to Epi.M or Epi.I in our culture collection were screened in amplicon sequencing data set, revealing correlation between culture-dependent and independent results. Genera such as *Arthrobacter, Staphylococcus, Priestia*, and *Bacillus* were predominantly found in mock treated plants, while the genera *Pseudomonas* and *Enterobacter*, as well as *Pantoea* correlated with the infection fraction. “Epi” = epiphytes. “Endo_W” = Endophytes after epiphyte wash. Absolute abundance after normalization: a threshold is set according to the sample that has the least sequence number, then the number of sequences set by this threshold are selected for analysis. * Indicates statistical significance (student’s t-test) p < 0.05. Figure 2A was created with Biorender.com

By analysing the amplicon sequencing data of the epiphytic and endophytic fractions and comparing the genus of these selected strains (**Dataset S4**), we confirmed that these strains can function as marker strains. Strikingly, as shown by the amplicon sequencing data, *Staphylococcus* and *Arthrobacter* genera were more abundant in both epiphytic and endophytic non-infected fractions compared to infected samples (**Fig. 2C**). *Bacillus* spp. and *Priestia* spp. were exclusively found in non-infected fractions and were more abundant in the epiphytic fraction (**Fig. 2D**). In contrast, the genus *Paenibacillus* was found in both non-infected and infected samples (**Fig. 2E**). Isolates of the genus *Enterobacter* and *Pseudomonas* were enriched in the epiphytic infected fraction (**Fig. 2F**).

A previous study showed that *Pantoea dispersa* and *Pantoea agglomerans* isolated from infected maize plants have a strong inhibitory activity against *U. maydis* and thus are crucial targets of the antimicrobial effector Ribo1^37^. While *P. dispersa* was not isolated in the present study, amplicon sequencing data revealed this genus as a strong marker for the endophytic fraction in infected leaves (**Fig. 2G**). We therefore included the two previously characterized strains in this study and subjected them to full genome sequencing (**Table S1**). This identified the previously 16sRNA-based classified *P. agglomerans* as the closely related *Enterobacter sichuanensis* (**Table S1**). Thus, we grouped *Pantoea dispersa* and *Enterobacter sichuanensis* in a community designated as “PdEs” for further experiments.

### Exploring the role of antimicrobial effectors in intermicrobial interactions of *U. maydis*

Next, we aimed to investigate a potential involvement of fungal proteins in the drastically shifted leaf microbiome after *U. maydis* infection. Antimicrobial effectors have previously been implicated in shaping leaf-associated microbial communities^24,56^. We therefore tested whether bacterial isolates recovered from maize leaves are sensitive to antimicrobial effectors produced by *U. maydis*. Given that Ribo1 has been previously functionally characterized in context with PdEs^37^, it was included as a reference effector in subsequent assays. Moreover, we investigated a possible role of a secreted GH25 lysozyme encoded by *U. maydis gh25* (gene ID: UMAG_02727). GH25 has been identified as an antimicrobial effector in the epiphytic yeast *Moesziomyces bullatus* ex *Albugo* (short*: MbA*), which is closely related to *U. maydis*. The biological function of GH25 in *U. maydis* is unknown, however, we previously found that GH25 antimicrobial activity is conserved between GH25 orthologs of *MbA* and *U. maydis*^38^. Thus, we aimed to explore if GH25 might act as an antimicrobial effector that contributes to *U. maydis* virulence. Re-visiting of publicly available RNA-seq datasets revealed similar expression dynamics for both *U. maydis gh25* and *ribo1*, with transcript levels peaking at early infection stages (0.5–1 dpi) and strongly decreasing at later time points (**Fig. 3A,B**)^57^. To assess a possible contribution of GH25 to *U. maydis* virulence, we generated a *gh25* knockout strain (SG200Δ*gh25*), as well as a strain constitutively overexpressing *gh25* (SG200_pOtef_*gh25*). In addition, a *gh25/ribo1* double-knockout mutant was generated for subsequent infection assays in maize. Similarly, to *ribo1,* deletion of *gh25* did not influence *U. maydis* virulence in controlled growth chamber conditions (**Fig. 3C1** for disease abundance and **Fig. 3C2** for disease index and statistical evaluation). Thus, the antimicrobial GH25 is not required for the pathogenic development of *U. maydis*. However, overexpression of *gh25* did slightly increase *U. maydis* virulence (see **Fig. S5A**) which is in contrast to *ribo1*, where overexpression lead to strong virulence reduction in *U. maydis* due to its cytotoxic activity to the host^37^.

**Figure 3:**
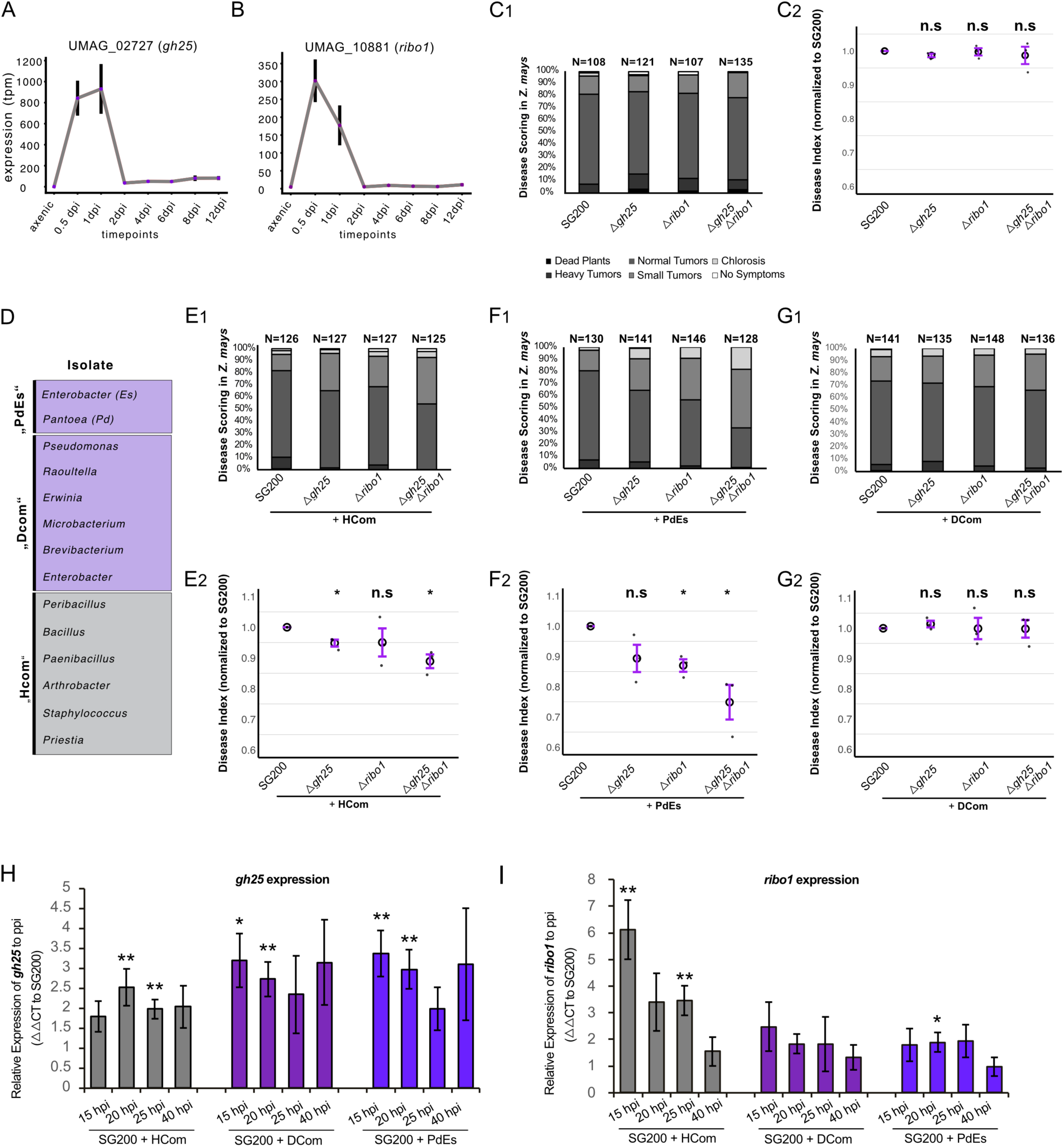
Role of *Ustilago maydis* antimicrobial effectors upon co-inoculation of maize plants bacterial communities. **A**) Expression pattern of UMAG_02727 (*Um_gh25*) and **B**) UMAG_10881 (*Um_ribo1*) – data retrieved from Lanver et al (2018) RNAseq. **C1**) Disease Scoring of effector knockout mutants and solopathogenic strain SG200, which serves as WT. Different disease classes are scored and represented by different colors. N indicates number of scored plants. **C2**) Relative Disease Index is calculated based on scored disease symptoms (normalized to SG200) and one disease index value was obtained per biological replicate; mutant values were normalized to the corresponding WT within each experiment (WT = 1). Statistical significance was assessed using a one-sided one-sample t-test against the reference value 1 (n = 3 biological replicates). **D**) Three different communities were selected based on culture-dependent and independent results: “HCom” = health-associated community, “DCom” = disease-associated community, and “PdEs” which including two previously described isolates targeted by the antimicrobial effector Ribo1^1^. **E-G**) Co-inoculation infection assay of effector knockout strains with HCom/DCom/PdEs E1-G1) Disease Scoring of effector knockout mutants and solopathogeneic strain SG200, which serves as WT. Co-inoculation with **E1**) HCom, **F1**) PdEs and **G1**) DCom. Different disease classes are scored and represented by different colors. N indicates number of scored plants. **E2-G2**) Relative Disease Index is calculated based on scored disease symptoms (normalized to SG200) and one disease index value was obtained per biological replicate, and mutant values were normalized to the corresponding WT within each experiment (WT = 1). Co-inoculation with **E2**) HCom, **F2**) PdEs and **G2**) DCom. Statistical significance was assessed using a one-sided one-sample t-test against the reference value of 1 (n = 3 biological replicates). Expression of **H**) *gh25* and **I**) *ribo1* were assessed after co-inoculation with either HCom, DCom or PdEs at 15, 20, 25 and 40 hpi. Quantitative PCR data were analyzed using the ΔΔCt method. Statistical analyses were performed on ΔΔCt values, while fold changes (2⁻ΔΔCt) are shown for visualization. For comparisons between treated and mock samples, two-sided one-sample t-tests against zero were applied using biological replicates as the unit of analysis. * Indicates statistical significance (p<0.0001:****, p<0.001:***, p<0.01:**, p<0.05:*, p>0.05: n.s.). Error bars show SDE.

### Health-associated (HCom) and disease-associated (DCom) communities influence the virulence of *U. maydis* differently, depending on the presence of the antimicrobial effector GH25

Subsequently, we tested for a potential role of *gh25* for plant infection in presence of distinct bacterial communities. For this, the members of the HCom, DCom, and PdEs communities (**Fig. 3D**) were used for co-inoculation experiments. For brevity, bacterial strains of HCom and DCom will hereafter be referred to by their genus name (e.g., *Priestia* in place of *Priestia megaterium*). As for the PdEs strains, *Pantoea dispersa* will be referred to as *Pantoea* (Pd) and *Enterobacter sichuanensis* as *Enterobacter* (Es).

We first tested whether these communities could influence virulence of the virulent *U. maydis* strain SG200, which was found true for PdEs and DCom, but not for the HCom (**Fig. S6A-C**). Next, we tested *U. maydis* strains SG200Δ*ribo1*^37^ and the newly generated SG200Δ*gh25* and SG200Δ*ribo1*Δ*gh25* strains in co-inoculation experiments with the three communities. This demonstrated impaired virulence of the deletion strains in presence of both HCom and PdEs, but not when co-inoculated with the DCom (**Fig. 3E-G**). As previously shown, deletion of *ribo1* led to reduced virulence upon co-inoculation with PdEs, while deletion of *gh25* led to reduced virulence in response to HCom. However, trends observed for both effector knockouts were similar. The effector double knockout strain showed the strongest reduction in its virulence compared to the progenitor strain SG200. Together, these results identify GH25 as an antimicrobial effector contributing to *U. maydis* virulence in presence of maize HCom bacteria. Furthermore, function of GH25 appears to be additive, though partially overlapping, with the cytotoxic Ribo1.

Since co-inoculation with different bacterial communities led to effector-dependent virulence reduction, we subsequently investigated how *gh25* and *ribo1* are regulated upon co-inoculation with bacterial communities. For this purpose, we collected samples after 15, 20, 25, and 40 hours post-inoculation (hpi). Analysis of gene expression by qRT-PCR revealed the highest expression levels for both *gh25* and *ribo1* at 15 hours post-infection (hpi), with a strong decline to almost zero by 40 hpi (**Fig. S5B**). Upon co-inoculation with HCom/DCom/PdEs, a significant induction of both *gh25* and *ribo1* gene expression could be observed (**Fig. 3H,I**). Interestingly, *ribo1* was upregulated strongly upon co-inoculation with HCom (6fold at 15hpi), but only weakly in response to DCom and PdEs, indicating a specific regulation in response to distinct bacterial communities (**Fig. 3I**). In contrast, *gh25* was induced in response to all tested communities, but more strongly by DCom/PdEs. Moreover, while *ribo1* expression dropped at 40 dpi even if co-inoculated with bacteria, *gh25* expression stayed slightly induced even at 40 hpi in response to DCom and PdEs (**Fig. 3H**).

### *U. maydis* inhibits HCom bacteria but not DCom bacteria via GH25

While performing co-inoculation experiments, we observed that mixing of *U. maydis* with DCom and PdEs led to an impaired filamentation of *U. maydis* on potato dextrose agar (PD)+charcoal plates (**Fig. 4C**). To identify single strains mediating this inhibition of fungal development *in vitro*, we performed filamentation assays in one-to-one interactions with all DCom/HCom/PdEs members. Comparing SG200 with SG200Δ*gh25* revealed that presence of most DCom and PdEs members reduces *U. maydis* filamentation of both SG200 and the mutant strain, with *Enterobacter*, *Pseudomonas*, and the PdEs strains having the strongest effect (**Fig. 4A,B, Fig. S7**). In contrast, HCom members had no negative effect on SG200 filamentation, while *Priestia* significantly inhibited filamentation of SG200Δ*gh25* (**Fig. 4B,D**). To directly measure the impact of GH25 protein activity, we performed confrontation assays of the bacterial strains with purified GH25 protein in a luciferase assay. Strikingly, all isolated HCom bacteria were targeted by GH25. Cell viability upon incubation with active GH25 dropped to nearly 0% for strains *Priestia*, *Arthrobacter*, and *Bacillus* (**Fig. 4F**). Only the *Staphylococcus* isolate was slightly inhibited on-plate but not impaired in cell viability, which is in line with previous results observed for *MbA_*GH25^38^. In sharp contrast, selected DCom as well as PdEs bacteria were not targeted by GH25 on-plate, with only minor reductions in cell viability observed for some strains (**Fig. 4F**).

**Figure 4:**
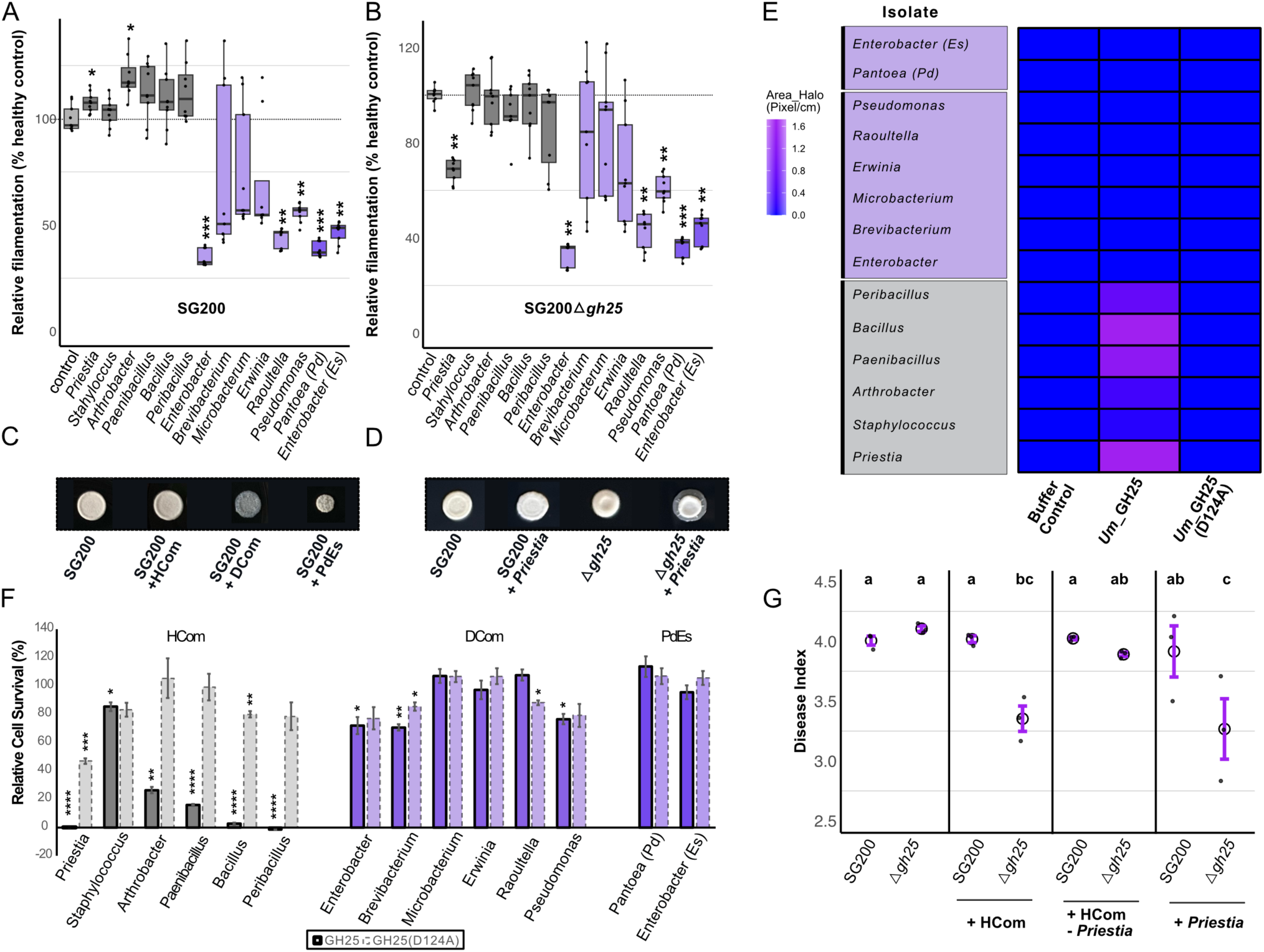
Bacterial communities influence *Ustilago maydis* filamentation virulence and filamentation, and are differentially targeted by GH25. **A,B**) Filamentation assay: For each biological replicate, mean pixel intensities were quantified for all spots. Technical replicates of the untreated control were averaged and used to normalize all measurements within the same biological replicate. Filamentation values were calculated as the ratio of control intensity to measured intensity, and expressed as percentage of healthy filamentation. Technical replicates were averaged prior to statistical analysis, and comparisons were performed on biological replicate means using Welch’s t-test. 100% line, representing the mean of the control treatment, is labelled by a dashed line. **C**) Filamentation of SG200 on PD + charcoal plates in mixture with DCom/HCom/PdEs. **D**) Filamentation of SG200 and *gh25* knockout mutant in mixture with *Priestia* **E)** Confrontation assay of all selected bacterial isolates with purified GH25 protein of *U. maydis*, including a buffer control and a protein containing a mutation in the active center (D124A) as negative control. Zone of inhibition was determined using ImageJ. **F**) Cell viability luciferase assay of all isolates to analyze % of bacterial cell survival upon treatment with Um_GH25, Um_GH25 (D124A) and buffer_control across 3 independent replicates. Luciferase counts were determined from three technical replicates per treatment. Data were averaged at the biological replicate level (n = 3 biological replicates per treatment). Statistical comparisons between treatments were performed using Welch’s two-sample t-test to account for unequal variances. * Indicates statistical significance (p<0.0001:****, p<0.001:***, p<0.01:**, p<0.05:*, p>0.05: n.s.). Error bars show standard error. **G**) Infection assay of SG200 and *gh25* knockout mutant in co-inoculation with either HCom, HCom – *Priestia*, or *Priestia* alone. For comparisons involving multiple treatments and/or mutants, unnormalized disease index values were used to allow direct statistical comparison among all groups, and differences were assessed using one-way ANOVA followed by appropriate post-hoc tests.

Based on the GH25-dependent filamentation phenotype together with the strong antimicrobial activity of GH25 towards *Priestia*, we asked whether this isolate might represent a key species in the observed virulence reduction of *U. maydis in planta*. Drop-out experiments and co-inoculation of maize with *U. maydis* and *Priestia* alone confirmed this hypothesis: the conditional virulence defect of the *U. maydis*Δ*gh25* mutant was lost upon removal of *Priestia* from the HCom (**Fig. 4G)**. Concordant, co-inoculation with *Priestia* alone could reproduce the previously observed phenotype for co-inoculation with the whole HCom (**Fig. 4G**). Thus, inhibition of *U. maydis*Δ*gh25* infection by the HCom can be attributed to the presence of *Priestia*.

### Cross-inhibition assays and functional profiling suggest distinct niche adaptation of HCom and DCom bacteria, reflecting *U. maydis* induced reprogramming of host physiology

The identified microbial communities show distinct patterns in their influence on *U. maydis* infection, as well as differential sensitivity to GH25. On the one hand, this reflects that *U. maydis* actively shapes the maize leaf microbiome during the early stages of infection. However, it is unlikely that the change in the bacterial community can be explained solely by the antimicrobial effect of *U. maydis*, particularly since it is unlikely that all bacteria are in direct spatial contact with the pathogen. To address this gap, we investigated two potential driving forces: (i) cross-inhibitory interactions within bacterial communities, and (ii) the role of nutrient acquisition in enabling communities such as HCom to colonize healthy maize leaves.

HCom bacteria might actively inhibit DCom bacteria and therefore more effectively colonize, allowing them to outcompete DCom bacteria on healthy plants. However, by performing cross-inhibition assays to assess the antagonistic capacity of individual strains, we found that DCom and PdEs bacteria exhibited a stronger antagonistic potential toward other strains, whereas most HCom bacteria were unable to grow when inoculated onto DCom bacterial lawns (**Fig. S8**). Consequently, the absence of DCom bacteria on healthy plants is unlikely to result from inhibition by HCom bacteria. Instead, we hypothesized that, besides the direct antagonistic activity of *U. maydis*, adaptation to distinct physiological environments in healthy versus infected plant promotes either HCom or DCom bacteria. It is important to mention that *U. maydis* induced tumors creates a strong metabolic sink, characterised by an increased uptake of carbohydrates and organic nitrogen^26,28,29^. We therefore re-examined published metabolomics analysis describing host metabolic reprogramming during *Ustilago maydis* infection of maize leaves^28^, which we summarized at the level of major metabolite classes relevant to nitrogen and carbon cycling (**Supplementary Table S2**). Tumor tissue is consistently characterized by increased pools of reduced organic nitrogen compounds, including free amino acids and polyamines, as well as elevated levels of soluble sugars, as reported by metabolomics and targeted sugar analyses of *U. maydis*–infected maize tissue^28,58,59^. In contrast, pools of inorganic nitrogen, including nitrate and ammonium, were reduced in tumor tissue relative to healthy leaves. Thus, we hypothesized that the niche to be colonized by bacterial communities fundamentally changes during infection, both structurally and metabolically, which might be reflected in the metabolic profile of each community.

To assess the effect of the altered metabolic environment on the growth of HCom and DCom, we constructed genome-scale metabolic models for both communities **(see Methods and Fig. 5A**). To recapitulate *in vivo* conditions, we performed stepwise growth simulations using a medium with shifting availability of key metabolites associated with either health- or tumor-associated conditions (**Fig. 5B, SI Appendix).** Starting from a metabolic environment that supports the growth of HCom and DCom nearly equally, we shifted the medium composition towards health-associated conditions by decreasing the availability of reduced organic nitrogen compounds (Gln, Asn, Arg, Putrescine) and soluble sugars (Glucose, Fructose) and increasing the availability of inorganic nitrogen (Nitrate, Ammonium). This shift led to a strong decrease in DCom growth which was far less pronounced in the HCom. On the contrary, an increase of reduced organic nitrogen compound and soluble sugar availability combined with a decrease in inorganic nitrogen availability, which represents tumor-associated conditions, enabled faster growth of the DCom compared to the HCom. Further analysis of shifts of individual metabolite subsets revealed that the availability of reduced organic nitrogen containing amino acids (Gln, Asn and Arg) was the main driver of the observed shift (**Fig. S10**).

**Figure 5:**
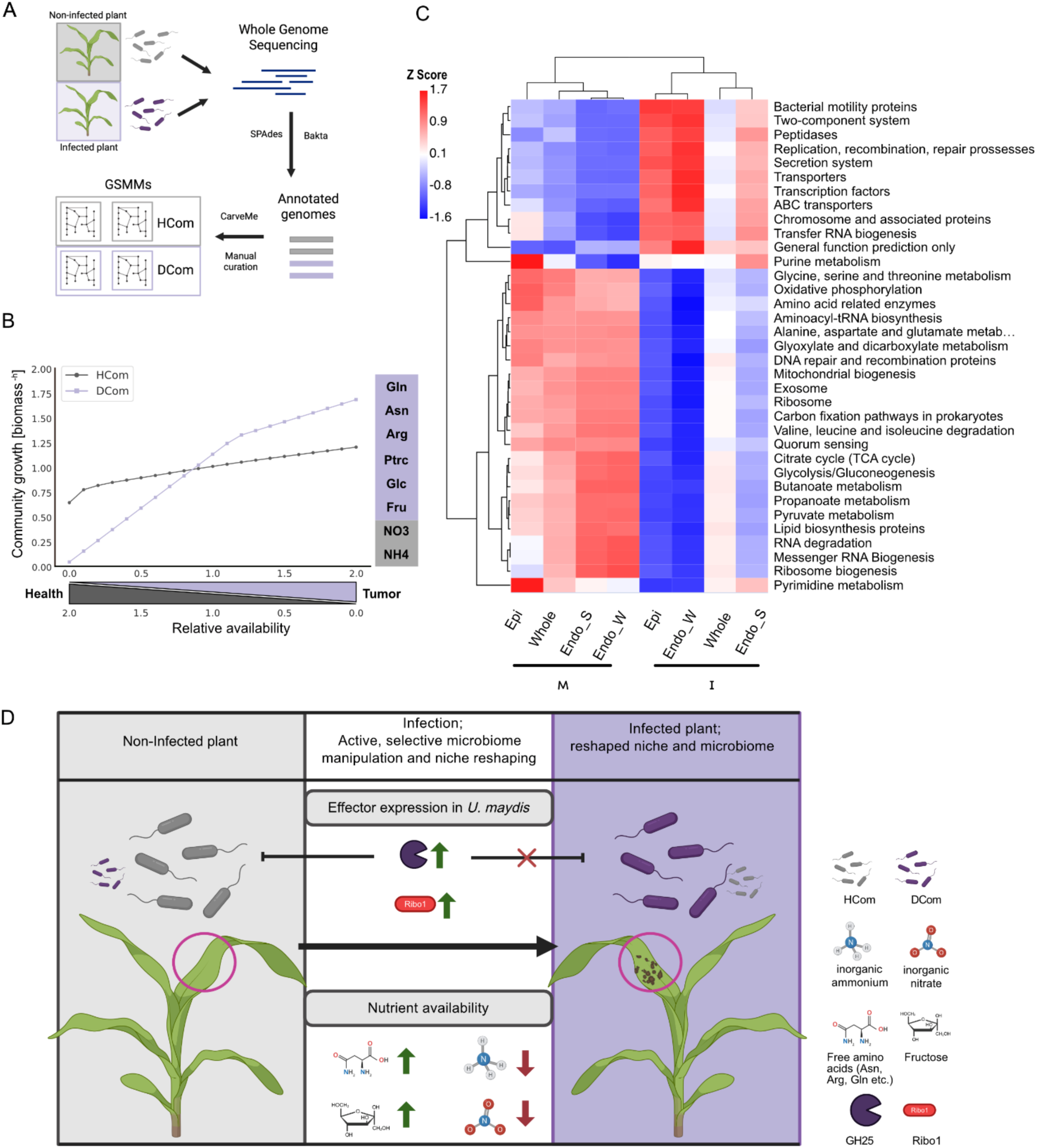
Additional factors influencing microbiome assembly. ***A*)** Overview of genome-scale metabolic model (GSMM) reconstruction. Annotated genome assemblies of all HCom and DCom members were used to reconstruct metabolic models of the individual members. Curated models were then combined into an HCom and a DCom community model. **B)** Simulating community growth under metabolic shifts into health and tumor-associated conditions by scaling the import fluxes of health- and tumor-associated metabolites. **C**) Tax4Fun Analysis Heatmap: functions are clustered to group those with similar compositions. Heatmap colors reflect Z-scores of row-wise relative abundances after standardization, so colors are comparable horizontally (same function across samples) but not vertically (different functions in the same sample). **D**) **Proposed Model: Two-step model of phyllosphere microbiome modulation during *U. maydis* infection.** In healthy plants, the phyllosphere is characterized by a microbiome dominated by healthy-associated taxa (HCom) and a nutrient environment with limited availability of soluble sugars and comparatively higher pools of inorganic nitrogen. During early infection, *U. maydis* actively and selectively modulates the phyllosphere microbiome through the secretion of antimicrobial effectors, including GH25 and Ribo1, which are expressed at this stage (expression data: this study and Lanver et al (2018)). These effectors directly target bacteria associated with mock-treated plants, leading to depletion of HCom members, while disease-associated taxa (DCom) are largely tolerant. At this stage, no consistent global shift in host nutrient availability is detected (Horst et al. 2010). At the tumor stage, microbiome modulation becomes predominantly indirect and niche-driven. Expression of early infection effectors is reduced, while tumor-associated effectors are induced and promote host reprogramming. Tumor development is associated with large-scale metabolic changes in infected tissue, including increased availability of soluble sugars and reduced pools of inorganic nitrogen. These host-derived changes restructure the phyllosphere niche, favoring copiotrophic and nitrate-reducing microbial taxa, and stabilizing a disease-associated microbiome via ecological selection rather than direct antimicrobial activity. Arrows indicate relative changes. Nutrient availability: comparison between healthy and infected seedlings. Solid arrows denote consistent global trends, while question marks indicate uncertainty. Arrows do not imply quantitative magnitude. Figure 5 A and D were created with Biorender.com

In line with the genome-scale metabolic modelling findings, TAX4Fun-based functional analysis revealed distinct differences in the predicted metabolic potential of microbial communities between non-infected and infected samples (**Fig. 5C**). Non-infected samples were enriched in pathways associated with central and anabolic metabolism, including carbon fixation, the citrate cycle (TCA), quorum sensing, and amino acid metabolism (glycine, serine, and threonine) (**Fig. 5C**). In contrast, infected samples showed a higher predicted abundances of pathways related to environmental interaction and resource acquisition, including secretion systems, peptidases, transporters, bacterial motility proteins, and transcription factors (**Fig. 5C**). FAPROTAX based functional annotation similarly shows that non-infected samples exhibited a higher absolute abundance of predicted functions related to nitrogen fixation, urea degradation (ureolysis), and methanol utilization, whereas infected samples showed a higher representation of predicted chemoheterotrophy and aerobic chemoheterotrophy, along with an increased abundance of functions related to nitrate reduction **(Fig. S9)**. Overall, the functional profiles differed markedly between non-infected and infected conditions.

Based on the combined results of microbiome profiling, effector assays, metabolic modelling, functional prediction, and a re-evaluation of published metabolomics data, we propose a model summarising the known interactions between *U. maydis*, the maize phyllosphere microbiome, and the host’s metabolic state (**Fig. 5D**). The model illustrates the differences in community composition between non-infected and infected leaves, as well as the selective activity of *U. maydis* antimicrobial effectors against specific bacterial taxa during the early stages of infection. Additionally, the model summarises the distinct nutrient landscapes inferred for non-infected and infected leaves. Therefore, we propose that the antimicrobial activity of *U. maydis* drives a niche adaptation involving the altered metabolic environment it induces in the plant.

## Discussion

Plant infection by filamentous fungal pathogens can lead to drastic changes in the microbiome structure of the host^24^. Here, we investigated the microbiome modulation in infection sites of *U. maydis*, representing an extreme and underexplored environment, to study microbial community assembly and the molecular mechanisms underlying this process. Infected maize leaves revealed drastic changes in microbiome composition in both epiphytic and endophytic communities, resulting in a reduction of alpha diversity. We found that the secreted antimicrobial effector UmGH25 plays a key role in the colonisation success of *U. maydis* in its host by driving a significant shift in the composition of the phyllosphere microbial community, thereby indicating directional adaptation through targeted microbial restructuring.

The observed reduction of bacterial diversity upon pathogen infection is similar to what has been previously demonstrated for other pathogens, such as *Fusarium oxysprorum* in *A. thaliana* roots and soybean roots^60,61^. However, a significant increase upon *U. maydis* infection was found for the *Pseudomonas and Enterobacter* species (**Fig. 1C**). In wheat, a strong enrichment of *Pseudomonas* was observed during *Fusarium graminearum* infection^62^. These *Pseudomonas* isolates were found to counteract pH changes induced by the pathogen through the production of organic acids. Another report focused on the foliar disease of *Pseudostelleria heterophylla*, which attracts *Pseudomonas* strains that could disrupt the production of fusaric acid by *Fusarium oxysporum* through the emission of volatile organic compounds^63^. Hence, enrichment of *Pseudomonas* upon pathogen infection might be partially attributed to active plant recruitment. Similarly, *Enterobacter* was previously isolated from necrotic rice blast lesions after *Magnaporthe oryzae* infection, indicating that this genus might be associated with infection sites^64^. Since disease-induced microbiome shifts can have many causes, the enrichment of specific genera could be due to various factors. For example, drivers can be direct recruitment by the plant or manipulation by the pathogen. However, community shifts can also be caused by niche manipulation, i.e. shifts in the abundance of specific metabolites^20^.

We found that bacterial strains isolated from infected leaves exhibited antagonistic potential against *U. maydis in vitro*. Thus, one could hypothesise host-driven recruitment of these strains. However, co-inoculation with DCom and PdEs bacteria had only a marginal impact on the virulence of *U. maydis*, suggesting additional factors to drive community shift. In recent work, Garrido-Sanz and Keel (2025) demonstrated that seed-borne rhizosphere bacteria (SbRB) act as dominant early colonizers of the wheat rhizosphere, often outcompeting or pre-empting native soil microbes during the initial stages of community assembly^65^. They found that SbRB (including *Pantoea, Priestia, Pseudomonas* and *Paenibacillus* genera), which were enriched in the early plant rhizosphere, carry specialised metabolic traits (e.g. saccharide transport and metabolism, and vitamin biosynthesis) that provide an advantage when utilising the simple and complex carbohydrates secreted by the host, as well as facilitating cross-feeding interactions with partner taxa. In this study, we found genera as *Priestia, Arthrobacter* and *Bacillus* to be enriched in non-infected plants. These HCom members were found to be antagonized by *U. maydis* through its antimicrobial effector GH25. Thus, one could hypothesize that these strains are prevalent in non-infected plants and appear to compete directly with *U. maydis*. Their ability to arrive early and acquire nutrients may help them establish a protective microbial landscape before pathogen invasion. By contrast, GH25 does not target DCom/PdEs, which suggests that *U. maydis* may not naturally encounter these strains on the leaf surface. Interestingly, *gh25* gene expression was induced in response to both bacterial communities, while *ribo1* expression was only upregulated in response to HCom. This might further hint towards HCom being the prevalent target, since *ribo1* expression appears to be more delicate for *U. maydis*, as its overexpression strongly reduced virulence^37^, while constitutive overexpression of *gh25* had no negative effect on *U. maydis*.

Growth simulations using genome-scale metabolic community models further support the hypothesis that host metabolic reprogramming during infection selectively favors disease-associated microbial communities. Under simulated health-associated conditions, characterized by reduced soluble sugars and reduced organic nitrogen compounds but increased inorganic nitrogen availability, DCom growth was suppressed, whereas HCom remained comparatively stable. In contrast, simulated leaf-tumor conditions enriched in soluble sugars and reduced organic nitrogen compounds promoted substantially faster DCom growth relative to HCom. Additional simulations further indicated that this competitive advantage was primarily driven by increased availability of free amino acids, whereas sugars alone had no significant effect **(Fig. S10A-D**). Moreover, under unconstrained nutrient conditions, DCom exhibited inherently faster growth, consistent with a community structured around metabolically opportunistic taxa adapted to nutrient-rich environments (**Fig. S10E**). These findings suggest that *U. maydis*-induced metabolic reprogramming may not merely alter microbiome composition indirectly but instead creates a nutrient environment that promotes opportunistic bacterial assemblages.

FAPROTAX-based functional inference and TAX4Fun predictions indicate that infection is linked to a change in the predicted metabolic potential of the microbial community. Samples from uninfected plants showed an enrichment of functions related to nitrogen fixation, urea degradation and methanol utilisation as well as an enrichment of pathways related to central and anabolic metabolism, including carbon fixation, quorum sensing and amino acid metabolism. These functions are commonly associated with nutrient recycling and metabolic versatility under stable environmental conditions^66^. These traits may reflect a stable, health-associated community structured around internal nitrogen turnover and the utilisation of specific carbon sources. By contrast, infected samples showed a greater prevalence of chemoheterotrophy, aerobic chemoheterotrophy, nitrate reduction, as well as pathways related to transport, secretion systems, motility, and environmental responsiveness. Other studies have reported similar patterns in FAPROTAX-based analyses following fungal infection, where diseased rhizomes showed an enrichment of opportunistic bacteria (e.g. *Klebsiella* and *Enterobacter*) and functional shifts identified by FAPROTAX, including an enrichment of functions associated with heterotrophy and nitrogen cycling in disease-affected samples^67^.

While predictive functional approaches should be interpreted cautiously, their agreement with genome-scale metabolic simulations strengthens the model that infection-associated microbiomes are metabolically adapted to the host metabolic reprogramming and niche restructuring during *U. maydis* infection^26,28^. Recent work has further demonstrated that this metabolic rewiring is actively driven by fungal effectors that reprogram host transcriptional networks controlling carbon allocation, nitrogen metabolism, and cell proliferation^68,69^. Within this altered biochemical landscape, the increased presence of nitrate-reducing and chemoheterotrophic species in infected plants appears to be an ecologically logical occurrence. By contrast, the association of healthy plants with nitrogen fixation indicates a comparatively nitrogen-limited phyllosphere in the absence of infection, where costly processes such as nitrogen recycling offer a selective advantage. Furthermore, the increased dominance observed in infected communities, coupled with reduced Shannon diversity, indicates that infection favours a limited number of taxa that competitively exclude others.

However, in contrast to a scenario of strong deterministic filtering, iCAMP^40^ analysis indicates that community assembly in infected tissues is primarily governed by dispersal-related and stochastic processes, with only a minor contribution of homogeneous selection. This suggests that infection disrupts host-mediated filtering and opens ecological niches that can be readily colonized by incoming taxa. Under such conditions, community composition is shaped less by consistent selective pressures and more by colonization dynamics, priority effects, and local stochasticity, resulting in variable but dominance-structured communities. Future studies could examine the potential recruitment of bacterial communities in areas not directly infected by the pathogen, as well as identifying additional antimicrobial effectors with different expression patterns. This would improve our understanding of direct microbiome manipulation during *U. maydis* infection.

In summary, this work shows that infection by *U. maydis* drastically alters the composition of the host microbiome, decreasing bacterial diversity and potentially paving the way for opportunistic bacteria. To compete with the microbiome of healthy plants, *U. maydis* deploys the antimicrobial effectors GH25 and Ribo1. We propose a model in which manipulation of the host microbiome occurs through two main processes: (i) colonization, which is characterized by the secretion of antimicrobial effectors that directly target and reduce the abundance of early colonisers, and (ii) established infection, which alters the leaf surface niche. The tumor-as-sink effect then results in an increased presence of opportunistic strains. Together, the results shown in this study provide a first step towards a mechanistic understanding of how *U. maydis* infection reshapes the maize phyllosphere microbiome, through a combination of effector-mediated antimicrobial activity and host metabolic reprogramming.

## Material and Methods

### Growth conditions for microbial strains

*U. maydis* strains were grown in liquid YEPS light medium at 28°C on a rotary shaker (200 rpm) and maintained on potato dextrose agar (PD) plates. Isolated bacterial strains (**Dataset S3** for all isolated bacteria) were grown in dYT or Luria Bertani (LB) liquid media at 28°C overnight on a rotary shaker (200 rpm). *Pichia pastoris* KM71H-OCH was used for recombinant protein expression as described in Eitzen et al. (2021)^70^.

### Plant growth conditions

For culture-dependent and culture-independent collection approaches for the analysis of the *Z. mays* microbiome, grains of maize (*Z. mays*, cv. Golden Bantam) were sown in organic growth soil topped with approximately 10% (v/v) of homogenized soil collected from a field plot where maize plants were cultivated, located at the coordinates 50.92548° N, 6.93594° E.. Plants were inoculated using the local compatible wild strains Um_45^71^ and Um_48^71^, according to Kämper et al. (2004)^72^. Plants were grown outdoor at the site with the coordinates 50.925392° N, 6.936483° E in September 2023 for the generation of bacterial culture collection from the maize phyllosphere. The experiment was replicated in August 2025 for the culture-independent approach, where amplicon sequencing of the bacterial endophytes and epiphytes, as well as both fractions combined, was performed.

### Culture collection of the maize phyllosphere

For the generation of the culture collection, 6 biological replicates per treatment (non-infected and infected with the wild compatible strains) were used. At 15 dpi, the infected region and its corresponding area in non-infected plants were sampled. A total of 5 leaf disks with 1.2 cm diameter from each biological replicate were used. Epiphytes were collected in 10 mM MgCl_2_ supplemented with 0.01% Tween20. Endophytes were collected through surface sterilization of the leaves by dipping them in 70% ethanol until they are fully coated then swiftly passing them through a flame of a Bunsen burner until the ethanol is dried, avoiding damaging/burning the leaves. Leaves were ground and suspended in 10 mM MgCl2 and dilutions was plated on LB agar and incubated for 2-3 weeks to allow the growth of slow-growing bacteria. The identification of the isolates was done by amplifying and sequencing the hypervariable regions of the 16S rRNA gene **(Dataset S5)**.

### Amplicon sequencing of the maize phyllosphere

For amplicon sequencing i.e., the culture-independent approach and analysis of the *Z. mays* microbiome, growth and sampling conditions are as described above. Four biological replicates per treatment (non-infected and infected with the wild compatible strains) were used. Processing of the samples took place directly after sampling (detailed methodology **in SI Appendix**). For sampling the whole microbiota, leaf sections were only washed with sterile Milli-Q water to remove airborne contaminants and loosely attached particles. As for the epiphytic fraction, we established modifications to the isolation protocol described by Li et al. (2022)^73^ to account for the susceptibility of maize seedling leaves to shearing forces. In short, epiphytes were dislodged from the leaf surface and released into PBS buffer (pH = 7.2) supplemented with 0.01% Tween20 via gentle sonication and shaking conditions, and subsequently collected in 0.2 μm cellulose nitrate membranes (25 mm diameter) (Whatman™, Cytiva; Merck, Darmstadt, Germany).

The endophytic fraction was isolated in 2 different methods in order to ensure full and consistent representation of the endophytes. The first method entailed the use of leaf sections that were used to isolate the epiphytes i.e., the resulting isolated microbiota would consist of the entire endophytic fraction, plus the epiphytes that remained tightly attached to the leaves after the epiphyte isolation. The second method of collecting the endophytes was done as described in the “Culture collection of the maize phyllosphere” section. DNA extraction from the leaf material was done using an SDS lysis buffer followed by a phenol/chloroform/isoamyl alcohol purification protocol, RNase digestion, and protein precipitation. As for the epiphytic fraction collected in the cellulose nitrate membrane, DNA extraction was performed following a custom protocol combining chemical lysis by an SDS buffer and mechanical lysis by bead beating, followed by a phenol/chloroform/isoamyl alcohol purification protocol^74^.

Amplicon libraries were prepared from pooled PCR products, end-repaired, A-tailed and ligated with Illumina adapters by Novogene (Munich, Germany). The libraries were sequenced on an Illumina NovaSeq 6000 platform using a paired-end strategy **(SI Appendix)**.

### Colony forming units and leaf imprint analysis

Fresh plant material was weighed and processed to quantify culturable bacteria from different plant-associated microbial fractions. To this end, the wash buffer containing the epiphyte fraction, and ground material resuspended in PBS buffer containing the endophyte fractions and whole microbiota (as described in the previous sections) were used. Bacterial suspensions from all fractions were subjected to serial tenfold dilutions in sterile 10 mM MgCl2 buffer. From each dilution, 5 µL were plated in technical triplicates onto LB agar medium using the drop plate method and incubated until colony formation was visible CFU values were back-calculated to the original suspension and normalized to plant fresh weight according to the following formula:

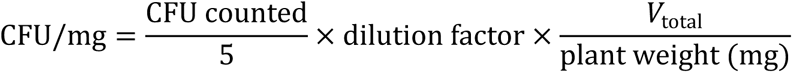

where 𝑉_total_ denotes the total volume of the respective bacterial suspension (e.g. 120 mL for wash samples). Results are reported as CFU per mg plant material (CFU/mg). Log10 transformation was applied where indicated.

### Microbial confrontation assays

For bacterial confrontation assays, bacterial cultures were diluted to OD_600_ 0.6 in 10 mM MgCl2 and 100 µl culture were spread on PD plates. 20 µl of GH25 protein (0.8µg/µl) was applied to a hole created in the centre of the plate (d=4mm). For bacterial cross-inhibition assays, square TSA plates were used. Bacteria were diluted to an OD_600_ of 0.8 in 10 mM MgCl2 and 250 µl of a lawn bacterium was spread. 5 µl of competitor bacteria were dropped on the plate. Plates were incubated for 2-4 days at 28°C. Pictures of the whole plate were taken with a ruler for reference. Zones of inhibition (Halos) were then measured using the size reference, evaluating them in ImageJ.

### Cell viability assays

To assess cell viability after GH25 treatment, a luciferase assay using the BacTiter-GloTM Microbial Cell Viability Assay (Promega, Madison, USA) was performed as described in Sorger et al. (2025)^38^. In short, each HCom, DCom and PdEs strain was grown overnight and then diluted to an OD_600_ of 0.1 in 10 mM phosphate buffer in 200 µl reactions. Active and mutated (D124A) *U. maydis* GH25 was added to a final concentration of 10 µM. A buffer-only control was included, and reaction was incubated for 1 hour/37°C. Protocol was performed according to the manufacturer’s instructions and relative survival was calculated by setting the buffer control to 100% and using the following formula: (“RLU_treated”/”RLU_buffer_control”)*100.

### Infection assays

The *U. maydis* transformation assay was performed by using protoplasts prepared according to Kämper (2004)^72^. Disease assays and co-inoculations for the solopathogenic strain of *U. maydis* SG200, SG200Δ*ribo1*, SG200Δ*gh25* and *SG200Δgh25Δribo1* strains were performed according to Ökmen et al.^37^. Each *U. maydis* cell suspension was injected exclusively into the juvenile stems of 6 to 7-d-old maize seedlings (GB) at the third leaf stage, via a syringe with needle (detailed infection protocol and statistical analysis approach in **SI Appendix)**.

### Filamentation assays

For filamentation assays, *U. maydis was* prepared as described in Ökmen et al.^37^; *U. maydis* cultures were mixed with a bacterial cultures (*U. maydis*: OD_600_ 0.6, Bacteria: OD_600_ 0.2) and 5µl were plated on PD + charcoal plates. Mean pixel intensities were quantified after two days using identical acquisition and analysis settings for all samples using Bio-Rad Image Lab software (6.0.1). Higher pixel intensity values corresponded to increased inhibition of filamentation in this assay. For each biological replicate, technical replicates of the untreated (low-inhibition) control were averaged to obtain a replicate-specific reference value. All measurements within the same biological replicate, including control samples, were normalized to this reference to account for inter-replicate variability. Filamentation was calculated as the ratio of control intensity to measured intensity and expressed as percentage of healthy filamentation (% of control), such that values close to 100% indicate uninhibited filamentation and lower values indicate increased inhibition. Technical replicates were averaged prior to statistical analysis. Statistical comparisons were performed on biological replicate means using Welch’s t-test. Data are presented as mean ± SD.

### Nucleic acid methods for gene expression analysis

For the analysis of gene expression of antimicrobial effectors, total RNA extraction from plant samples were performed with Trizol Reagent (Invitrogen, Karlsruhe, Germany) according to the manufacturer’s instructions followed by treatment with Turbo DNA-Free Kit (Ambion/Applied Biosystems; Darmstadt, Germany) to remove any DNA contamination in the extracted RNA. Synthesis of cDNA was performed using First Strand cDNA Synthesis Kit (Thermo Fisher Scientific, Darmstadt, Germany) according to recommended instruction starting with 3 µg RNA. Relative expression levels of marker genes were analyzed with GoTaq® qPCR Master Mix in a Bio-Rad iCycler system using the following program: 2 min at 95 °C followed by 45 cycles of 30 s at 95 °C, 30 s at 61 °C and 30 s at 72 °C. Primers used in qRT-PCR can be found in **Dataset S5**.

### Construction of plasmids

All plasmids were constructed using Gibson assembly (New England Biolabs; Frankfurt a.M., Germany). Plasmid Prep Kit (QIAGEN, Venlo, The Netherlands) was used for isolation of plasmid DNA from bacteria. GH25 active site mutants were created by using QuickChange XL Site-Directed Mutagenesis Kit from Agilent Technology according to the manufacturer’s instructions. *E. coli* transformation was performed using heat shock according to standard molecular biology methods^75^. Vector constructs and oligonucleotide sequences are mentioned in **Dataset S5**. Generated constructs were verified at the Eurofins sequencing facility (Germany).

### Fungal transformation and validation

The *Um*Gh25 overexpression constructs (**Dataset S5**) were introduced in *SG200* by ip integration, using protoplasts according to Kämper (2004)^72^. Generated strains were further analyzed by southern blot to validate the integration and copy number.

### Statistical analysis

All statistical analyses were performed using biological replicates as the unit of analysis. For CFU assays, values from technical replicates were averaged per plant, and plant-level means were log₁₀-transformed prior to analysis. Differences between treatments were assessed using Welch’s two-sample t-test to account for unequal variances. Disease symptoms of infection assays were scored for individual plants and summarized as a disease index per biological replicate. For single comparisons to the SG200 control, disease indices were normalized to SG200 (SG200 = 1) and analyzed using a one-sample t-test against the reference value of 1. For comparisons involving multiple treatments and/or mutants, non-normalized disease index values were used to allow direct statistical comparison among all groups, and differences were assessed using one-way ANOVA followed by appropriate post-hoc tests. Quantitative PCR data were analyzed using the ΔΔCt method. Statistical analyses were performed on ΔΔCt values, while fold changes (2⁻ΔΔCt) are shown for visualization. For comparisons between treated and mock samples, two-sided one-sample t-tests against zero were applied. Unless stated otherwise, all statistical tests were two-sided, and statistical significance was assessed using a threshold of *P* < 0.05.

### De novo whole genome sequencing (WGS) of HCom, DCom, and PdEs strains

The DNA of the bacterial strains of HCom, DCom, and PdEs was extracted (LGC Bioresearch technologies, distributed by Biozym, Hessisch Oldendorf, Germany). DNA samples were sent to Novogene facility (Munich, Germany) where libraries were prepared by DNA fragmentation, followed by end-repair, A-tailing, and ligation with Illumina adapters. The libraries were sequenced on Illumina NovaSeq X Plus platform using a paired-end strategy. Paired end sequencing results underwent quality control using the fastq toolbox for python 0.23.4^76^ and were subsequently used for genome assembly using the SPAdes De Novo Assembler 4.2.0^77^. The assembled genomes underwent quality control using checkm2 0.4^78^ and were annotated using Bakta 1.12.0^79^. Initial taxonomic assignment of all strains was performed using the Type (Strain) Genome Server (TYGS; DSMZ, Germany)^48–55^, which applies whole-genome-based phylogenomic analysis and digital DNA–DNA hybridization (dDDH) comparisons against curated type strain genomes for species-level classification. Taxonomic classification was subsequently refined and confirmed using GTDB-Tk 2.7.1^41–47^ with the classify workflow. Due to the substantial computational requirements of this workflow, including reliance on a large reference database and high memory usage (approximately 152 GB RAM), analyses were conducted on the ITCC high-performance computing (HPC) system RAMSES, utilizing 64 CPU cores and 160 GB of RAM.

### Metabolic network reconstruction and flux modelling

Genome-scale metabolic models for all 12 members of the HCom and DCom were reconstructed from the annotated genomes using CarveMe 1.6.6^80^. Models were curated following standard practices^81^ with particular focus to mass and charge balancing and removal of duplicate reactions. Model quality was evaluated using MEMOTE 0.17.0^82^ yielding an average overall quality score of ∼87%. Maximization of growth (using the biomass function) was set as the objective function for each individual microbe. All simulations were performed in Python 3.10 using COBRApy 0.30.0^83^.

Community models for the HCom and DCom were constructed using the Community() function in MICOM 0.37.1^84^, with individual strain models provided as input with predefined abundances based on the amplicon sequencing data of the maize phyllosphere. The community objective was defined as maximization of the summed growth rates of all members. Flux solutions were obtained using flux balance analysis (FBA)^85^. The growth medium was composed of primary metabolites identified in maize leaf extracts by HPLC^28^ supplemented with the combined output of the complete_community_medium() function of the HCom and DCom models (**SI Appendix**). Relative abundance of health- and tumor-associated metabolites were accounted for by applying a multiplicative factor to the uptake bounds of the respective metabolites of each individual member of the HCom and DCom; at 1, uptake bounds are unmodified and support near-equal growth. Shifts toward health were represented by decreasing tumor-associated metabolite uptake bounds (0.9–0.0) and increasing health-associated metabolite uptake bounds (1.1–2.0), with the reverse applied for tumor-associated conditions. Growth of both community models was simulated using the same medium composition and uptake bounds as modified for each simulation step.

## Data availability

All source data are provided in this paper and its supplementary information. Raw sequencing data is available online in the NCBI database under the following BioProject accession: PRJNA1441378. Associated Biosample and SRA records for both metagenomic and whole-genome sequencing datasets are available under the same BioProject. Assembled genome sequences are also available in GenBank under this BioProject, with each genome associated with a separate BioSample accession. The genome of *Raoultella ornithinolytica* is deposited in NCBI under the organism name *Klebsiella ornithinolytica*; in this paper, we refer to the species by the newer synonym *R. ornithinolytica.* All microbial strains are available on request from the corresponding author. Metabolic modeling medium composition and all code needed to reproduce the results of this study can be found at https://github.com/Toepfer-Lab/Maize_HCom_DCom

## Author contributions

GD, ZS and SD designed the study. ZS, SD, FR and BÖ performed experiments. ZS, SD, NT and GD evaluated the data. ZS and SD wrote the manuscript with input from all authors.

## Acknowledgements

This project has received from the Cluster of Excellence on Plant Sciences (CEPLAS) funding under Germany’s Excellence Strategy—EXC 2048/1—project ID: 390686111, the DFG priority program SPP2125 ‘DECRyPT’ and the Ministerium für Kultur und Wissenschaft des Landes Nordrhein-Westfalen (Ministry of Culture and Science of the State of North Rhine-Westphalia) (Profilbildung 20222–iHEAD—project number PB22-025A). We thank Dr. Janina Werner and Dr. Weiliang Zuo for their support in the experimental design. We thank Novogene for their support with data analysis and for their valuable collaboration throughout this project. We furthermore thank the ITCC (IT Center University of Cologne) for providing compute resources on the DFG-funded HPC (High Performance Computing) system RAMSES (Research Accelerator for Modeling and Simulation with Enhanced Security) as well as support (DFG funding number: INST 216/512-1 FUGG).

## Supplementary information

**Figure S1:** Overview of the culture collection and amplicon sequencing experiment workflow

**Figure S2:** Infection with *U. maydis* increases abundance of fast-growing bacteria

**Figure S3:** Infection with *U. maydis* shifts microbiome composition

**Figure S4:** Infection with *U. maydis* significantly increased the abundance of both epiphytes and endophytes in the phyllosphere

**Figure S5:** Expression of *gh25*/*ribo1* and the influence on *U. maydis* virulence

**Figure S6:** Co-inoculation of maize seedlings with *U. maydis* and bacterial communities

**Figure S7:** Charcoal filamentation assay plates

**Figure S8:** Cross-inhibition assay of all isolated strains from HCom/DCom and PdEs

**Figure S9:** FAPROTAX based functional analysis comparing Epi_Mock and Epi_Inf.

**Figure S10:** Effect of individual metabolite subsets on community growth

**Table S1:** Taxonomic assignment of the bacterial strains of the HCom, DCom, and PdEs communities as identified by GDTB-Tk (ANI based analysis) and TYGS (dDDH d4-based analysis)

**Table S2:** Summary of metabolite shifts associated with *U. maydis* tumor formation in maize leaves

**Table S3:** Summary of software and tools used for the analysis of the amplicon sequencing data.

**Dataset S1**: List of all ASVs identified in Amplicon Sequencing

**Dataset S2**: Alpha Diversity Values from Amplicon Sequencing

**Dataset S3**: Sequences and information of all isolated strains in culture collection

**Dataset S4**: Feature Table of each genus (absolute counts) from Amplicon Sequencing

**Dataset S5**: List of all oligonucleotides used in this study

## References

1. Ali, Q. et al. Power of plant microbiome: A sustainable approach for agricultural resilience. Plant Stress 14, 100681 (2024).

2. Mukherjee, A. et al. Unearthing the power of microbes as plant microbiome for sustainable agriculture. Microbiol. Res. 286, (2024).

3. Santos, L. F. & Olivares, F. L. Plant microbiome structure and benefits for sustainable agriculture. Curr. Plant Biol. 26, 100198 (2021).

4. Keppler, A. et al. Plant microbiota feedbacks through dose-responsive expression of general non-self response genes. Nat. Plants *2024 111* 11, 74–89 (2024).

5. Uribe-Acosta, M., Pascale, A., Zhou, J., Stringlis, I. A. & Pieterse, C. M. J. Roots: metabolic architects of beneficial microbiome assembly. Plant Physiol. 199, 349 (2025).

6. Wankhade, A., Wilkinson, E., Britt, D. W. & Kaundal, A. A Review of Plant–Microbe Interactions in the Rhizosphere and the Role of Root Exudates in Microbiome Engineering. Appl. Sci. 2025*, Vol.* 15, *Page* 7127 **15**, 7127 (2025).

7. Pascale, A., Proietti, S., Pantelides, I. S. & Stringlis, I. A. Modulation of the Root Microbiome by Plant Molecules: The Basis for Targeted Disease Suppression and Plant Growth Promotion. Front. Plant Sci. 10, 1741 (2020).

8. Santoyo, G. How plants recruit their microbiome? New insights into beneficial interactions. J. Adv. Res. 40, 45–58 (2022).

9. Jacoby, R. P., Koprivova, A. & Kopriva, S. Pinpointing secondary metabolites that shape the composition and function of the plant microbiome. J. Exp. Bot. 72, 57–69 (2021).

10. Vogel, C., Bodenhausen, N., Gruissem, W. & Vorholt, J. A. The Arabidopsis leaf transcriptome reveals distinct but also overlapping responses to colonization by phyllosphere commensals and pathogen infection with impact on plant health. New Phytol. 212, 192–207 (2016).

11. Compant, S., Samad, A., Faist, H. & Sessitsch, A. A review on the plant microbiome: Ecology, functions, and emerging trends in microbial application. J. Adv. Res. 19, 29–37 (2019).

12. Flor, H. H. & Pieterse, C. M. J. The Extended Plant Immune System. 10.1094/MPMI-10-25-0144-HH 38, 780–795 (2025).

13. Sommer, A. et al. A salicylic acid-associated plant-microbe interaction attracts beneficial Flavobacterium sp. to the Arabidopsis thaliana phyllosphere. Physiol. Plant. 176, e14483 (2024).

14. Korenblum, E. et al. Rhizosphere microbiome mediates systemic root metabolite exudation by root-to-root signaling. Proc. Natl. Acad. Sci. U. S. A. 117, 3874–3883 (2020).

15. Lebeis, S. L. et al. Salicylic acid modulates colonization of the root microbiome by specific bacterial taxa. Science *(80-.).* 349, 860–864 (2015).

16. Dastogeer, K. M. G., Yasuda, M. & Okazaki, S. Microbiome and pathobiome analyses reveal changes in community structure by foliar pathogen infection in rice. Front. Microbiol. 13, 949152 (2022).

17. Kuang, L. et al. Diseased-induced multifaceted variations in community assembly and functions of plant-associated microbiomes. Front. Microbiol. 14, 1141585 (2023).

18. Tie, Z. et al. Different responses of the rhizosphere microbiome to Verticillium dahliae infection in two cotton cultivars. Front. Microbiol. 14, 1229454 (2023).

19. Chao, S. et al. Exserohilum turcicum Alters Phyllosphere Microbiome Diversity and Functions—Implications for Plant Health Management. Microorganisms 13, 524 (2025).

20. Thomazella, D. P. T., Pereira, L. B. & Teixeira, P. J. P. L. Understanding microbiome shifts and their impacts on plant health during pathogen infections. Plant Physiol. 199, 498 (2025).

21. Berendsen, R. L., Pieterse, C. M. J. & Bakker, P. A. H. M. The rhizosphere microbiome and plant health. Trends Plant Sci. 17, 478–486 (2012).

22. Li, J., Wang, C., Liang, W. & Liu, S. Rhizosphere Microbiome: The Emerging Barrier in Plant-Pathogen Interactions. Front. Microbiol. 12, 772420 (2021).

23. Vogel, C. M., Potthoff, D. B., Schäfer, M., Barandun, N. & Vorholt, J. A. Protective role of the Arabidopsis leaf microbiota against a bacterial pathogen. Nat. Microbiol. *2021 612* 6, 1537–1548 (2021).

24. Mesny, F., Bauer, M., Zhu, J. & Thomma, B. P. H. J. Meddling with the microbiota: Fungal tricks to infect plant hosts. Curr. Opin. Plant Biol. 82, 102622 (2024).

25. Mesny, F., et al. Plant-associated fungi co-opt ancient antimicrobials for host manipulation. bioRxiv 2024.01.04.574150 (2025) doi:10.1101/2024.01.04.574150.

26. Doehlemann, G. et al. Reprogramming a maize plant: transcriptional and metabolic changes induced by the fungal biotroph Ustilago maydis. Plant J. 56, 181–195 (2008).

27. Matei, A. et al. How to make a tumour: cell type specific dissection of Ustilago maydis-induced tumour development in maize leaves. New Phytol. 217, 1681–1695 (2018).

28. Horst, R. J. et al. Ustilago maydis infection strongly alters organic nitrogen allocation in maize and stimulates productivity of systemic source leaves. Plant Physiol. 152, 293–308 (2010).

29. Kretschmer, M., Croll, D. & Kronstad, J. W. Maize susceptibility to Ustilago maydis is influenced by genetic and chemical perturbation of carbohydrate allocation. Mol. Plant Pathol. 18, 1222–1237 (2017).

30. Basse, C. W. & Steinberg, G. Ustilago maydis, model system for analysis of the molecular basis of fungal pathogenicity. Mol. Plant Pathol. 5, 83–92 (2004).

31. Villajuana-Bonequi, M. et al. Cell type specific transcriptional reprogramming of maize leaves during Ustilago maydis induced tumor formation. Sci. Reports 2019 91 9, 10227-(2019).

32. Lee, K., Pan, J. J. & May, G. Endophytic Fusarium verticillioides reduces disease severity caused by Ustilago maydis on maize. FEMS Microbiol. Lett. 299, 31–37 (2009).

33. Srivastava, D. A., Harris, R., Breuer, G. & Levy, M. Secretion-Based Modes of Action of Biocontrol Agents with a Focus on Pseudozyma aphidis. Plants 10, 210 (2021).

34. Harris, R. & Levy, M. Pseudozyma aphidis Extracts Possess Bioactive Properties That Suppress Plant Pathogens and Enhance Resistance in Tomato Plants. Plant Pathol. 74, 2184–2197 (2025).

35. Hewald, S. et al. Identification of a gene cluster for biosynthesis of mannosylerythritol lipids in the basidiomycetous fungus Ustilago maydis. Appl. Environ. Microbiol. 72, 5469–5477 (2006).

36. Teichmann, B., Linne, U., Hewald, S., Marahiel, M. A. & Bölker, M. A biosynthetic gene cluster for a secreted cellobiose lipid with antifungal activity from Ustilago maydis. Mol. Microbiol. (2007) doi:10.1111/j.1365-2958.2007.05941.x.

37. Ökmen, B. et al. A conserved extracellular Ribo1 with broad-spectrum cytotoxic activity enables smut fungi to compete with host-associated bacteria. New Phytol. 240, 1976–1989 (2023).

38. Sorger, Z., et al. GH25 lysozyme mediates tripartite interkingdom interactions and microbial competition on the plant leaf surface. Proc. Natl. Acad. Sci. 122, e2510124122 (2025).

39. Flores-Nunez, V. M. & Stukenbrock, E. H. The impact of filamentous plant pathogens on the host microbiota. BMC Biol. 22, 175 (2024).

40. Ning, D. et al. A quantitative framework reveals ecological drivers of grassland microbial community assembly in response to warming. Nat. Commun. *2020* 111 11, 4717-(2020).

41. Chaumeil, P. A., Mussig, A. J., Hugenholtz, P. & Parks, D. H. GTDB-Tk v2: memory friendly classification with the genome taxonomy database. Bioinformatics 38, 5315–5316 (2022).

42. Parks, D. H. et al. GTDB: an ongoing census of bacterial and archaeal diversity through a phylogenetically consistent, rank normalized and complete genome-based taxonomy. Nucleic Acids Res. 50, D785–D794 (2022).

43. Matsen, F. A., Kodner, R. B. & Armbrust, E. V. pplacer: linear time maximum-likelihood and Bayesian phylogenetic placement of sequences onto a fixed reference tree. BMC Bioinformatics 11, 538 (2010).

44. Shaw, J. & Yu, Y. W. Fast and robust metagenomic sequence comparison through sparse chaining with skani. Nat. Methods 2023 2011 20, 1661–1665 (2023).

45. Hyatt, D. et al. Prodigal: Prokaryotic gene recognition and translation initiation site identification. BMC Bioinformatics 11, 119-(2010).

46. Price, M. N., Dehal, P. S. & Arkin, A. P. FastTree 2 – Approximately Maximum-Likelihood Trees for Large Alignments. PLoS One 5, e9490 (2010).

47. Eddy, S. R. Accelerated Profile HMM Searches. PLOS Comput. Biol. 7, e1002195 (2011).

48. Meier-Kolthoff, J. P. & Göker, M. TYGS is an automated high-throughput platform for state-of-the-art genome-based taxonomy. Nat. Commun. *2019* 101 10, 2182-(2019).

49. Meier-Kolthoff, J. P., Carbasse, J. S., Peinado-Olarte, R. L. & Göker, M. TYGS and LPSN: a database tandem for fast and reliable genome-based classification and nomenclature of prokaryotes. Nucleic Acids Res. 50, D801–D807 (2022).

50. Freese, H. M., Meier-Kolthoff, J. P., Sardà Carbasse, J., Afolayan, A. O. & Göker, M. TYGS and LPSN in 2025: a Global Core Biodata Resource for genome-based classification and nomenclature of prokaryotes within DSMZ Digital Diversity. Nucleic Acids Res. 54, D884–D891 (2026).

51. Meier-Kolthoff, J. P., Auch, A. F., Klenk, H. P. & Göker, M. Genome sequence-based species delimitation with confidence intervals and improved distance functions. BMC Bioinformatics 14, (2013).

52. Lefort, V., Desper, R. & Gascuel, O. FastME 2.0: A Comprehensive, Accurate, and Fast Distance-Based Phylogeny Inference Program. Mol. Biol. Evol. 32, 2798–2800 (2015).

53. Kreft, L., Botzki, A., Coppens, F., Vandepoele, K. & Van Bel, M. PhyD3: a phylogenetic tree viewer with extended phyloXML support for functional genomics data visualization. Bioinformatics 33, 2946–2947 (2017).

54. Meier-Kolthoff, J. P. et al. Complete genome sequence of DSM 30083T, the type strain (U5/41T) of Escherichia coli, and a proposal for delineating subspecies in microbial taxonomy. Stand. Genomic Sci. 9, 2- (2014).

55. Farris, J. S. Estimating Phylogenetic Trees from Distance Matrices. 10.1086/282802 106, 645–668 (1972).

56. Snelders, N. C., Rovenich, H. & Thomma, B. P. H. J. Microbiota manipulation through the secretion of effector proteins is fundamental to the wealth of lifestyles in the fungal kingdom. FEMS Microbiol. Rev. 46, (2022).

57. Lanver, D. et al. The Biotrophic Development of Ustilago maydis Studied by RNA-Seq Analysis. Plant Cell 30, 300–323 (2018).

58. Sosso, D. et al. Sugar Partitioning between Ustilago maydis and Its Host Zea mays L during Infection. Plant Physiol. 179, 1373–1385 (2019).

59. Wahl, R., Wippel, K., Goos, S., Kämper, J. & Sauer, N. A Novel High-Affinity Sucrose Transporter Is Required for Virulence of the Plant Pathogen Ustilago maydis. PLOS Biol. 8, e1000303 (2010).

60. Kudjordjie, E. N. et al. Fusarium oxysporum Disrupts Microbiome-Metabolome Networks in Arabidopsis thaliana Roots. Microbiol. Spectr. 10, (2022).

61. Chang, H. X., Noel, Z. A. & Chilvers, M. I. A β-lactamase gene of Fusarium oxysporum alters the rhizosphere microbiota of soybean. Plant J. 106, 1588–1604 (2021).

62. Xu, Y. et al. Keystone Pseudomonas species in the wheat phyllosphere microbiome mitigate Fusarium head blight by altering host pH. Cell Host Microbe 0, (2025).

63. Yuan, Q. S. et al. Pathogen-driven Pseudomonas reshaped the phyllosphere microbiome in combination with Pseudostellaria heterophylla foliar disease resistance via the release of volatile organic compounds. Environ. Microbiome 19, 61 (2024).

64. Sahu, K. P. et al. Rice Blast Lesions: an Unexplored Phyllosphere Microhabitat for Novel Antagonistic Bacterial Species Against Magnaporthe oryzae. Microb. Ecol. 81, 731–745 (2021).

65. Garrido-Sanz, D. & Keel, C. Seed-borne bacteria drive wheat rhizosphere microbiome assembly via niche partitioning and facilitation. Nat. Microbiol. 2025 105 10, 1130–1144 (2025).

66. Fierer, N. et al. Comparative metagenomic, phylogenetic and physiological analyses of soil microbial communities across nitrogen gradients. ISME J. 2012 65 6, 1007–1017 (2011).

67. Wang, W. et al. Metabolome-driven microbiome assembly determining the health of ginger crop (Zingiber officinale L. Roscoe) against rhizome rot. Microbiome 12, 167 (2024).

68. Zuo, W., Depotter, J. R. L., Stolze, S. C., Nakagami, H. & Doehlemann, G. A transcriptional activator effector of Ustilago maydis regulates hyperplasia in maize during pathogen-induced tumor formation. Nat. Commun. 2023 141 14, 6722- (2023).

69. Lee, Y. J. et al. Ustilago maydis disrupts carbohydrate signaling networks to induce hypertrophy in host cells. bioRxiv 2024.11.22.624849 (2024) doi:10.1101/2024.11.22.624849.

70. Eitzen, K., Sengupta, P., Kroll, S., Kemen, E. & Doehlemann, G. A fungal member of the Arabidopsis thaliana phyllosphere antagonizes Albugo laibachii via a GH25 lysozyme. Elife 10, 1 (2021).

71. Depotter, J. R. L. et al. Effectors with Different Gears: Divergence of Ustilago maydis Effector Genes Is Associated with Their Temporal Expression Pattern during Plant Infection. J. Fungi 2021*, Vol.* 7, *Page 16* **7**, 16 (2020).

72. Kämper, J. A PCR-based system for highly efficient generation of gene replacement mutants in Ustilago maydis. Mol. Genet. Genomics (2004) doi:10.1007/s00438-003-0962-8.

73. Li, P. D. et al. The phyllosphere microbiome shifts toward combating melanose pathogen. Microbiome 10, 56- (2022).

74. Agler, M. T. et al. Microbial Hub Taxa Link Host and Abiotic Factors to Plant Microbiome Variation. PLoS Biol. 14, (2016).

75. Sambrook, J., Fritsch, E. F. & Maniatis, T. Molecular cloning: a laboratory manual. (Cold Spring Harbor Laboratory Press, 1989).

76. Chen, S. Ultrafast one-pass FASTQ data preprocessing, quality control, and deduplication using fastp. iMeta 2, e107 (2023).

77. Prjibelski, A., Antipov, D., Meleshko, D., Lapidus, A. & Korobeynikov, A. Using SPAdes De Novo Assembler. Curr. Protoc. Bioinforma. 70, e102 (2020).

78. Chklovski, A., Parks, D. H., Woodcroft, B. J. & Tyson, G. W. CheckM2: a rapid, scalable and accurate tool for assessing microbial genome quality using machine learning. Nat. Methods *2023 208* 20, 1203–1212 (2023).

79. Schwengers, O. et al. Bakta: Rapid and standardized annotation of bacterial genomes via alignment-free sequence identification. *Microb*. Genomics 7, 000685 (2021).

80. Machado, D., Andrejev, S., Tramontano, M. & Patil, K. R. Fast automated reconstruction of genome-scale metabolic models for microbial species and communities. Nucleic Acids Res. 46, 7542–7553 (2018).

81. Thiele, I. & Palsson, B. A protocol for generating a high-quality genome-scale metabolic reconstruction. Nat. Protoc. *2010 51* 5, 93–121 (2010).

82. Lieven, C. et al. MEMOTE for standardized genome-scale metabolic model testing. Nat. Biotechnol. *2020 383* 38, 272–276 (2020).

83. Ebrahim, A., Lerman, J. A., Palsson, B. O. & Hyduke, D. R. COBRApy: COnstraints-Based Reconstruction and Analysis for Python. BMC Syst. Biol. 7, 74- (2013).

84. Diener, C., Gibbons, S. M. & Resendis-Antonio, O. MICOM: Metagenome-Scale Modeling To Infer Metabolic Interactions in the Gut Microbiota. mSystems 5, (2020).

85. Orth, J. D., Thiele, I. & Palsson, B. O. What is flux balance analysis? Nat. Biotechnol. 2010 283 28, 245–248 (2010).

