## Supplementary information for "*Ustilago maydis* infection reshapes the maize phyllosphere microbiome through antimicrobial effectors and host metabolic reprogramming"

[*] contributed equally

**This file includes:**

Figures S1 to S10

Supplementary Table S1 (taxonomic assignments of HCom, DCom, and PdEs strains), S2 (metagenome revisiting) and S3 (summary of software and tools used for analysis of amplicon sequencing)

Supporting text: Material and Methods

SI References

**Other supporting materials for this manuscript include the following:**

Dataset S1: List of all ASVs identified in Amplicon Sequencing

Dataset S2: Alpha Diversity Values from Amplicon Sequencing

Dataset S3: Sequences and information of all isolated strains in culture collection

Dataset S4: Feature Table of each genus (absolute counts) from Amplicon Sequencing

Dataset S5: List of all oligonucleotides used in this study

#### Supplementary Figures

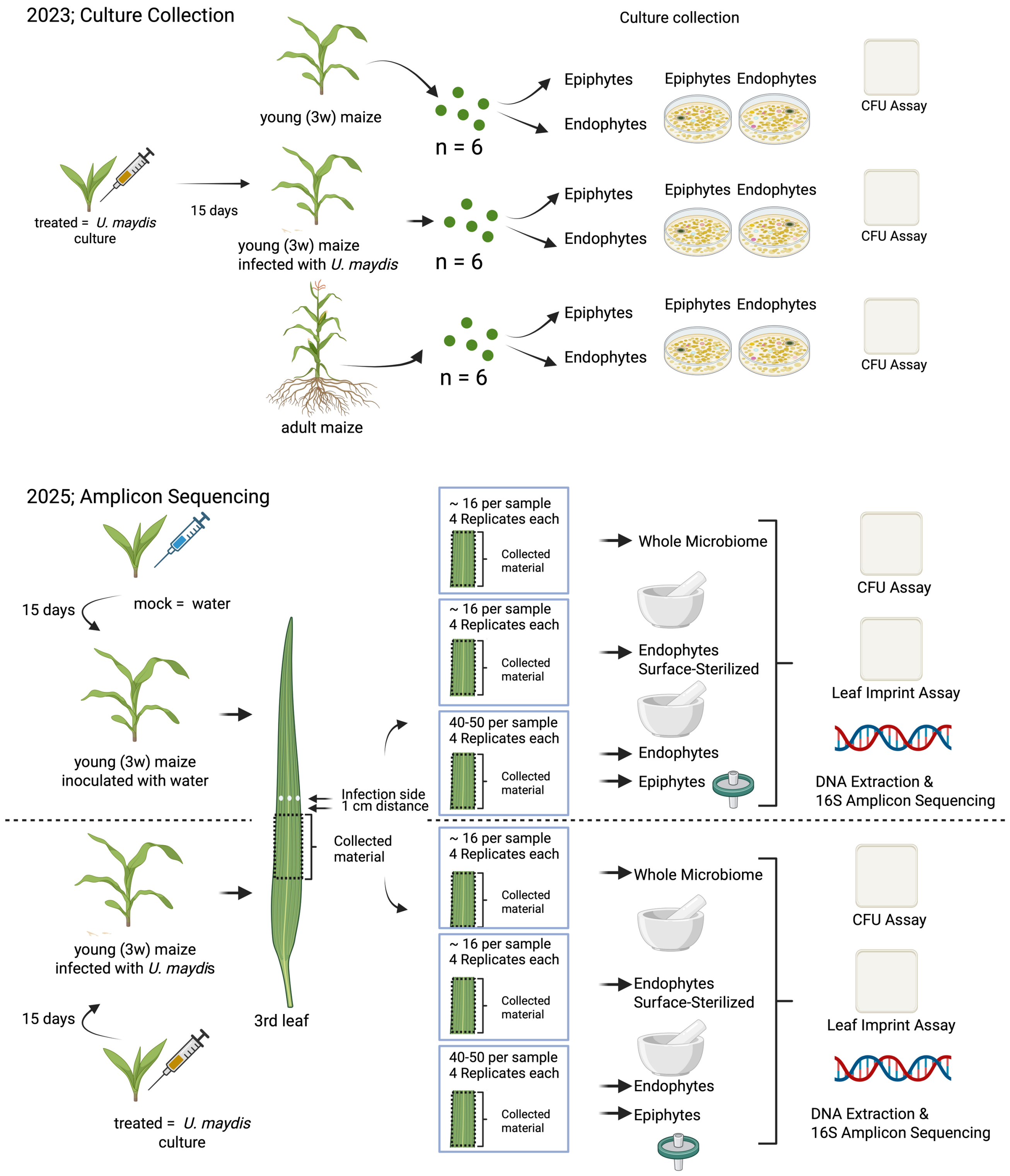


**Fig. S1: Overview of the culture collection and amplicon sequencing experiment workflow.** A culture collection was generated in 2023, by isolating bacteria from maize seedlings (var. Golden Bantam) seedlings infected with local compatible *U. maydis* strains. At 15 dpi, samples were collected from infected non-infected seedlings. In addition, leaf material was also collected from adult ky21 maize plants. Leaf disks were taken from infection sites (and corresponding areas in non-infected plants) and processed to isolate epiphytic and endophytic fractions. CFU assays were performed to investigate differences in bacterial colonization. In 2025, amplicon sequencing was performed. Plants were inoculated with either water (mock) or *U. maydis*, and samples were collected at 15 dpi from the tumor tissue region of the leaves. Four different fractions were isolated: epiphytic fraction, endophytic fraction after epiphyte wash (Endo_W), endophytic fraction after surface sterilization (Endo_S), as well as a whole microbiome fraction, where no separation occurred. A leaf imprint test and colony forming units (CFU) assay were performed before sample processing for amplicon sequencing. Created with Biorender.com

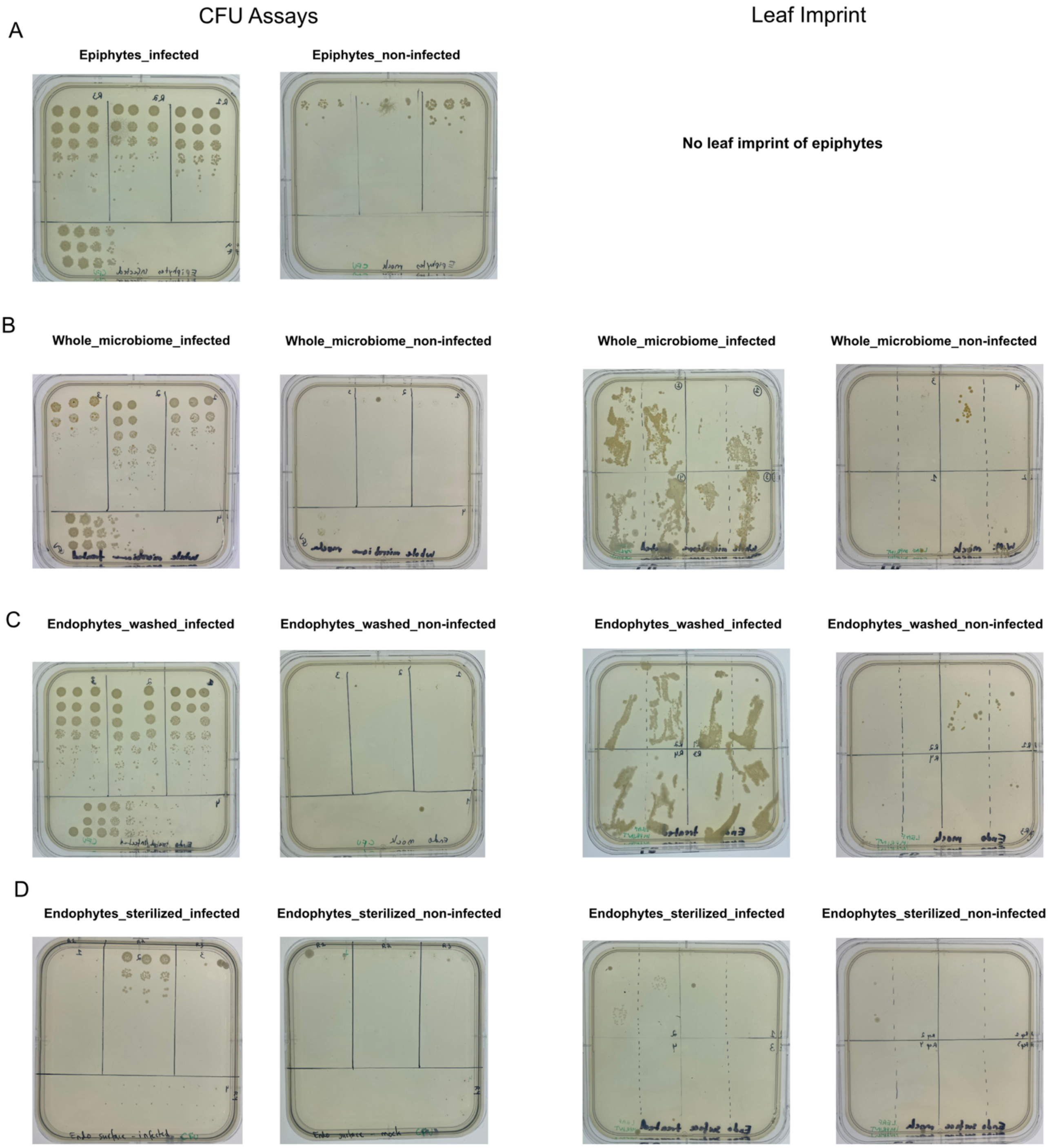


**Fig. S2: Infection with *U. maydis* increases abundance of fast-growing bacteria.** Colony forming unit and leaf imprint assay were performed alongside the amplicon sequencing approach. For entophytic samples, leaf material was taken, weighed and ground, whereas for the epiphytic samples, weight of total plant material was taken and part of the wash solution was used for CFU assay. Dilution series was performed and plated on LB plates. Growth of strains was assessed by counting the amount of colony forming units and calculating the amount of CFU (Log10(CFU/ml)).

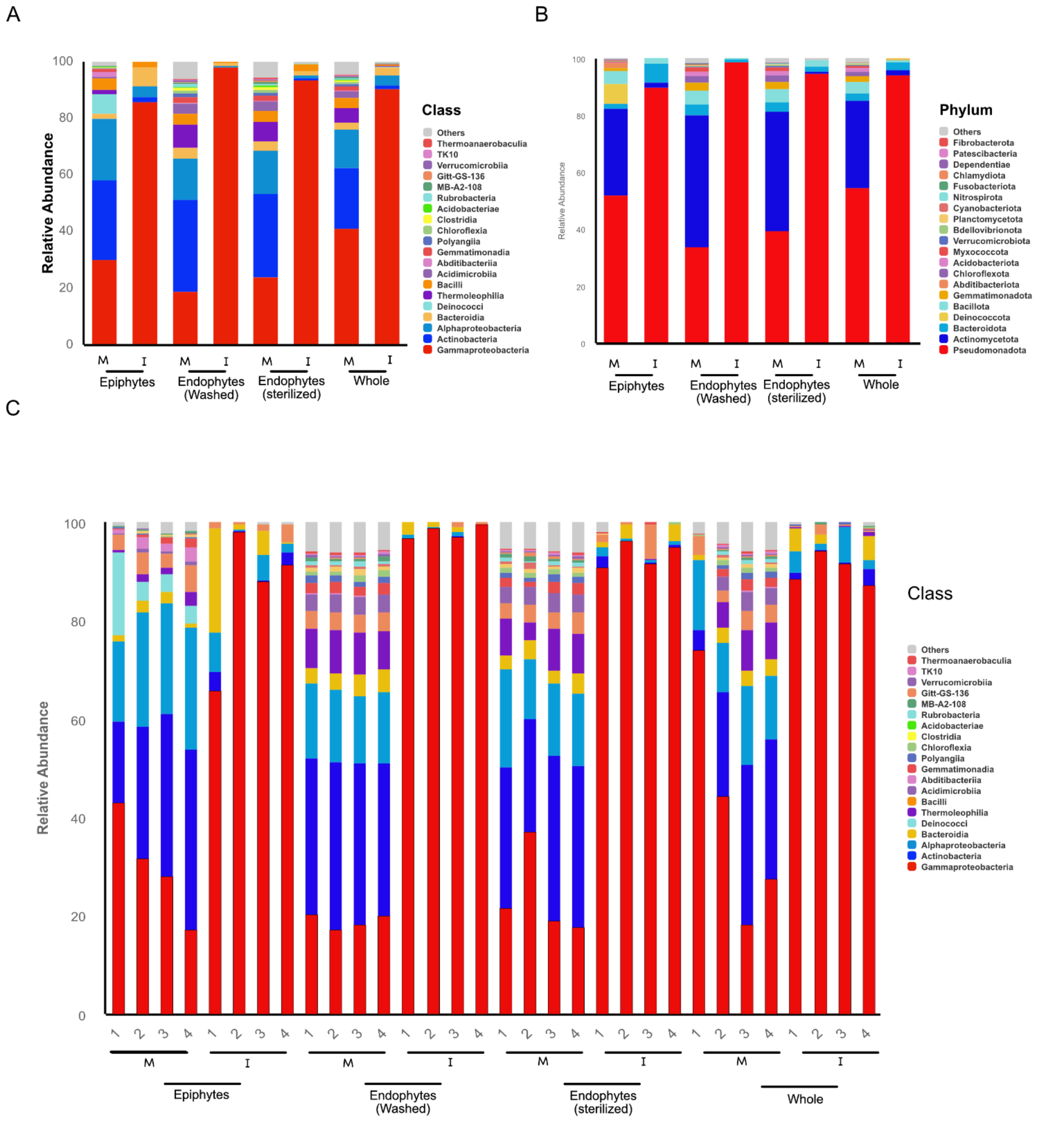


**Fig. S3: Infection with *U. maydis* shifts microbiome composition.** Relative abundances of main groups revealed a shift in the microbiome after *U. maydis* infected the plant. The top 20 taxa of each group at the taxonomic rank **A**) "Class" and **B**) "Phylum" were selected to form a distribution histogram of the relative abundance of these taxa. This allows the visualization of the taxa with a higher abundance and their proportion**. C**) The top 20 taxa of each sample at the taxonomic rank "class" were selected to form a distribution histogram of the relative abundance of these taxa.

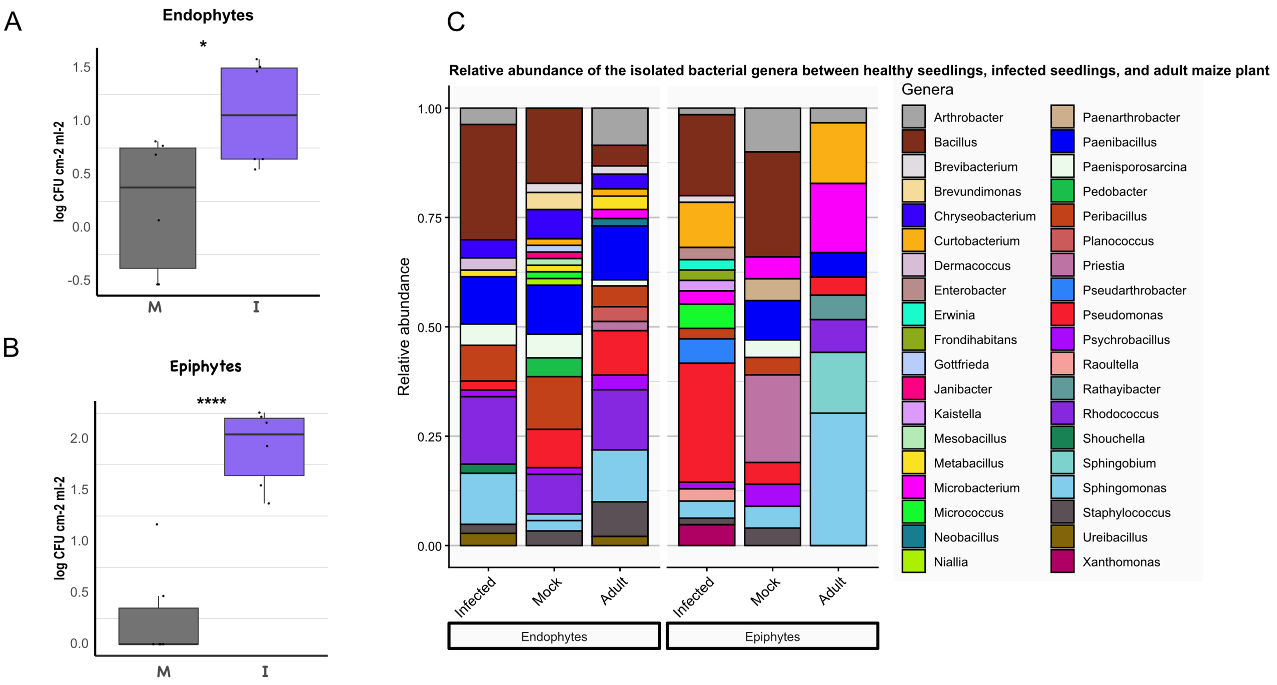


**Fig. S4: Infection with *U. maydis* significantly increased the abundance of both epiphytes and endophytes in the phyllosphere.** Colony forming units (CFU) assay was performed on samples taken for culture-dependent approach. Leaf disks were taken, grinded and a dilution series performed and plated to LB plates. Growth of strains was assessed by counting the amount of colony forming units and calculating the amount of CFU $ml^2cm^2$. **A**) comparison of abundance of epiphytes in mock and *U. maydis* infected plants. **B**) comparison of relative abundance of bacterial genera in non-infected seedlings, *U. maydis* infected seedlings, and healthy adult maize plants. **C**) comparison of abundance of endophytes in mock treated seedlings and mature plants. * Indicates statistical significance (Welch's t-test) $p < 0.05$.

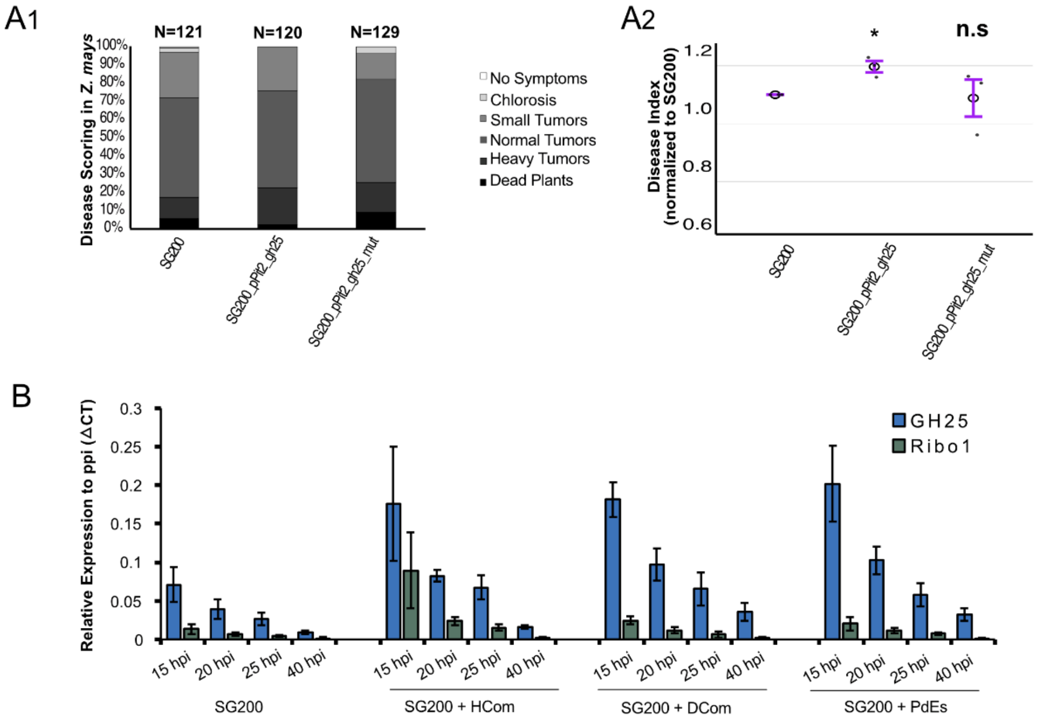


**Fig. S5: Expression of *gh25*/*ribo1* and the influence on *U. maydis* virulence**. **A**$_1$) Disease Scoring of GH25 effector gene overexpression strain and pathogenic strain SG200, which serves as WT. A mutant overexpressing active *gh25* under the pPit2 promotor, as well as a mutated version (D124A) was included. Different disease classes are scored and represented by different colors. N indicates number of scored plants. **A**$_2$) Relative Disease Index is calculated based on scored disease symptoms (normalized to SG200) and one disease index value was obtained per biological replicate; mutant values were normalized to the corresponding WT within each experiment (WT = 1). Statistical significance was assessed using a one-sided one-sample t-test against the reference value 1 (n = 3 biological replicates). **B**) Expression of *gh25* and *ribo1* were assessed after co-inoculation with either HCom, DCom or PdEs, as well as single inoculation (only SG200) at 15, 20, 25 and 40 hpi. Quantitative PCR data were analyzed using the ΔCt method.

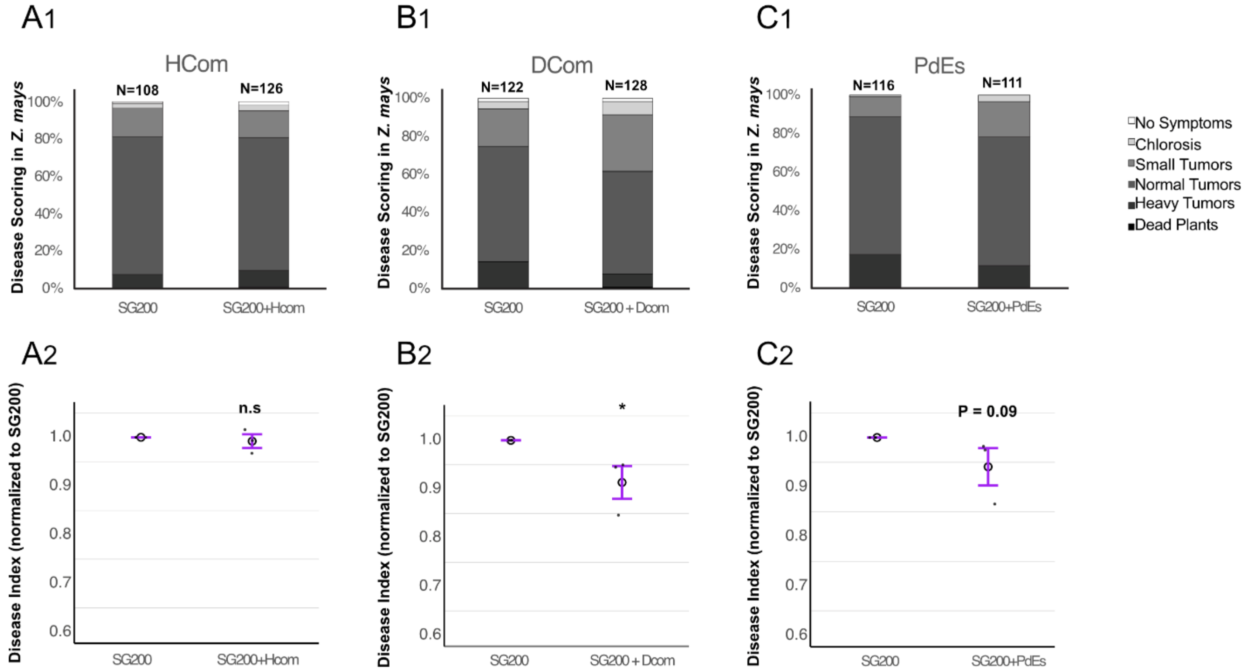


**Fig. S6: Co-inoculation of maize seedlings with U. maydis and bacterial communities. $A_1$ - $C_1$)** Disease scoring of *U. maydis* strain SG200, which serves as WT, alone and in co-inoculations with bacterial communities. Different disease classes are scored and represented by different colors. N indicates number of scored plants. **$A_2$ - $C_2$)** Relative Disease Index is calculated based on scored disease symptoms (normalized to SG200) and one disease index value was obtained per biological replicate; mutant values were normalized to the corresponding WT within each experiment (WT = 1). * Indicates statistical significance (p<0.0001:****, p<0.001:***, p<0.01:**, p<0.05:*, p>0.05: n.s.).

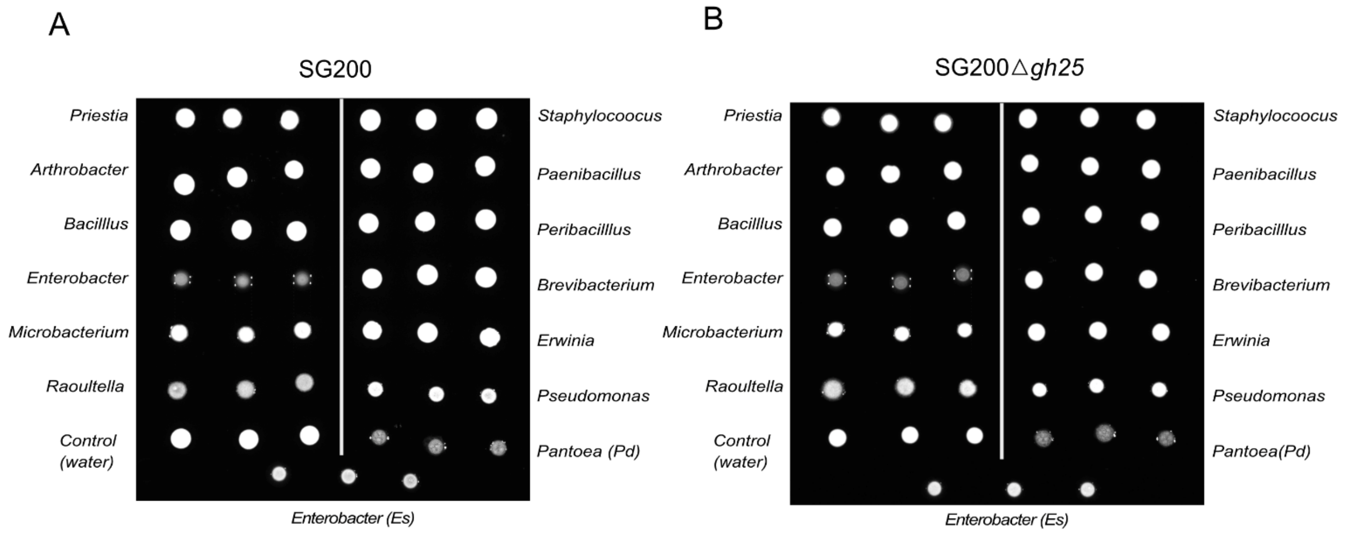


**Fig. S7: Charcoal filamentation assay plates.** Filamentation assay of *U. maydis* in response to different bacterial isolates. Strains were grown overnight, washed and resuspended in sterile water. Then, *U. maydis* and bacterial cultures were mixed to reach a final $OD_{600}$of 0.8 for *U. maydis* and 0.2 for bacterial competitor **A**) SG200 confronted with HCom/DCom/PdEs bacteria. **B**) Strain SG200Δ*gh25* confronted with HCom/DCom/PdEs bacteria. Mean Value (Intensity) was determined using ChemiDoc.

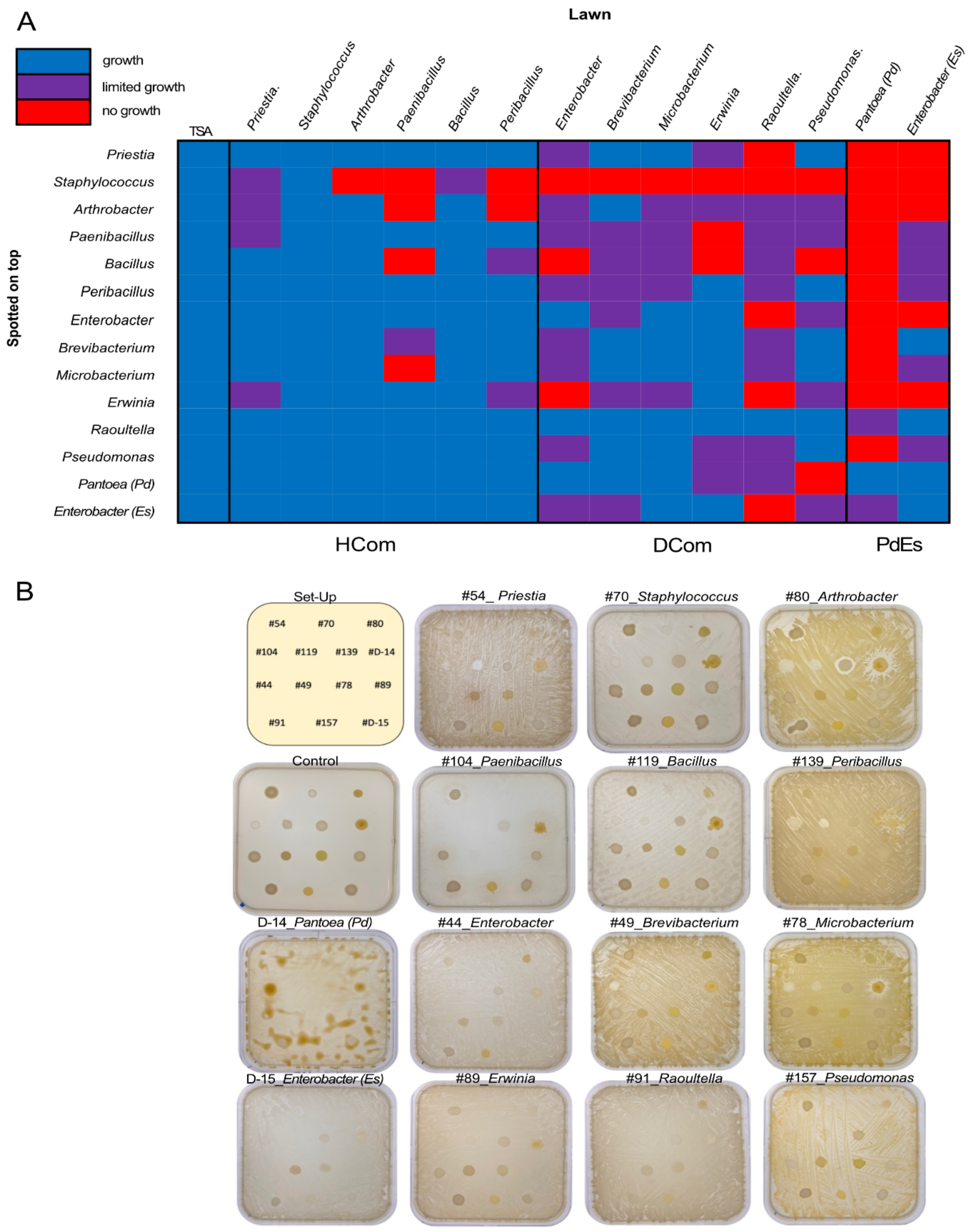


**Fig. S8: Cross-inhibition assay of all isolated strains from HCom/DCom and PdEs. A**) Interaction matrix and **B**) images of confrontation assays performed on TSA plates. 200 µl lawn bacterium was plated and 5µl treatment bacterium was dropped. Growth of treated bacteria was assessed after 2 days and scored in three categories: normal growth, weak growth and no growth. This experiment has been repeated twice, leading to reproducible results.

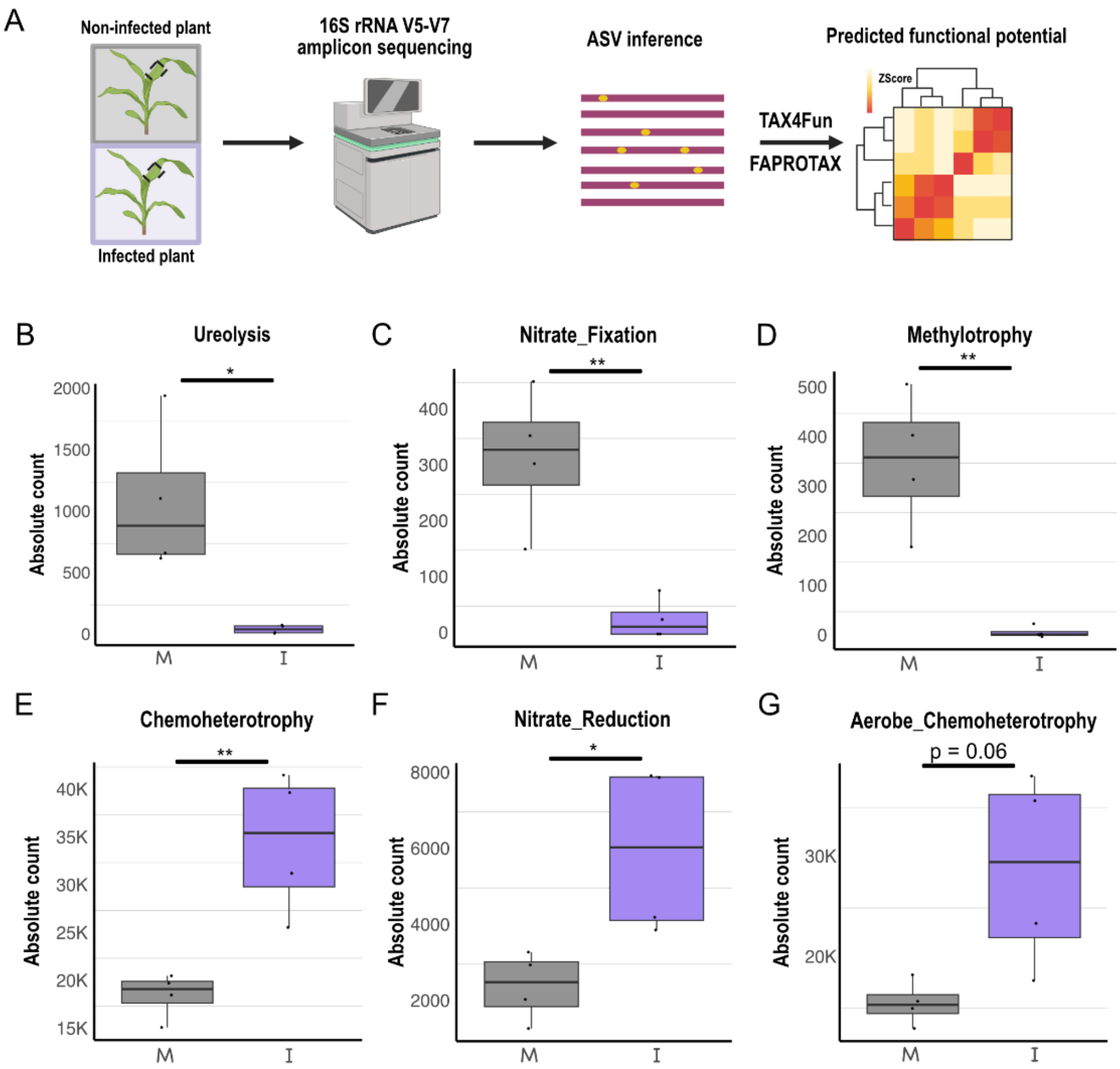


**Fig. S9: FAPROTAX based functional analysis comparing Epi_Mock and Epi_Inf. A**) Schematic overview of the experimental to functional prediction workflow. **B-G**) FAPROTAX based functional analysis comparing Epi_Mock and Epi_Inf. Absolute count is shown. FAPROTAX software has built a functional classification database based on species information. The current version, based on reference evidence, contains more than 80 functional classifications of carbon, nitrogen, phosphorus, sulfur and other elements, animal and plant pathogens, methane generation, and fermentation, covering more than 4600 different prokaryotic species, and has a good prediction effect on the biochemical cycle process of environmental samples. Each boxplot represents a different functional classification where differences between mock and infected were observed: **A**) Ureolysis, **B**) Nitrate_Fixation, **C**) Methylotrophy, **D**) Nitrate_Reduction, **E**) Aerobe_Cheromheterotrophy and **F**) Chemoheterotrophy. Differences between treatments were assessed using Welch's two-sample t-test to account for unequal variances. * Indicates statistical significance (p<0.0001:****, p<0.001:***, p<0.01:**, p<0.05:*, p>0.05: n.s.). Error bars show standard error. Fig. S9A was created with Biorender.com

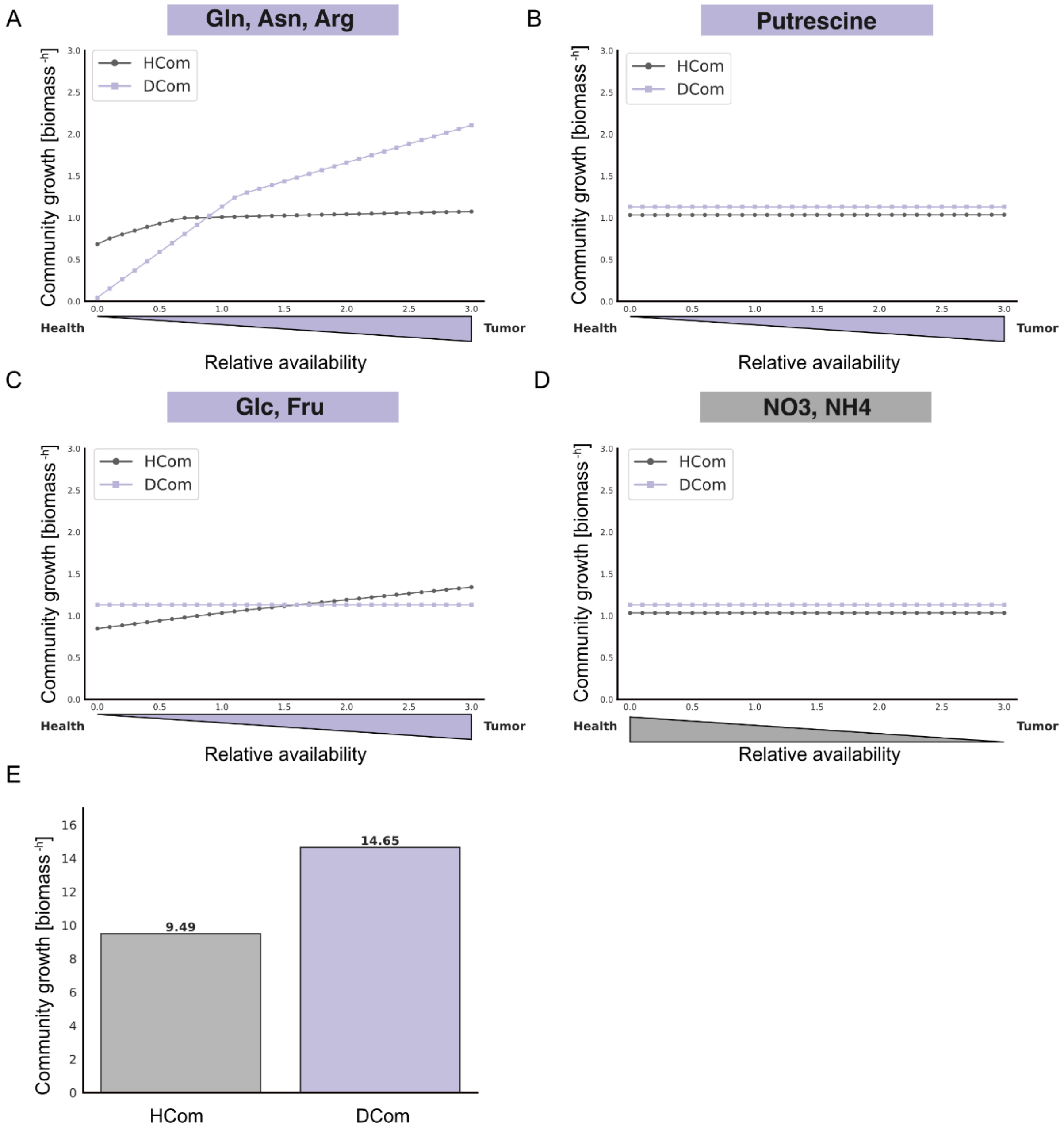


**Fig. S10**: **A-D**) **Effect of individual metabolite subsets on community growth**. Uptake bounds of **A**) [Gln, Asn, Arg], **B**) [Ptrc], **C**) [Glc, Fru] and **D**) [$NO_3$, $NH_4$] were independently scaled from 0 to 3 (step size 0.1), without applying changes to other metabolite uptake bounds. Growth rates of HCom and DCom were determined by flux balance analysis for each condition. **E**) **Growth under non-limiting conditions**. Growth-constraining metabolites were identified by reducing individual uptake bounds to 10% and selecting those causing >5% growth reduction. Uptake bounds of all constraining metabolites were subsequently set to non-limiting levels (1000), and growth of HCom and DCom was recalculated by flux balance analysis. Bars show resulting community growth rates.

#### Supplementary Material and Methods

##### Culture collection of the maize phyllosphere

For the generation of the culture collection, 6 biological replicates per treatment (infected and non-infected with the wild compatible strains) were used. At 15 dpi, the infected region and its corresponding area in the non-infected plants were sampled. A total of 5 leaf disks with 1.2 cm diameter from each biological replicate were used. Leaf disks were gently washed with 5 ml of sterile water for 10 seconds to remove loosely attached particles. Epiphytes were collected by shaking the leaf disks with 3 ml of 10 mM $MgCl_2$ supplemented with 0.01% Tween20, for approximately 2 minutes. Epiphytes were collected in the buffer. Endophytes were collected through surface sterilization of the leaves by dipping them in 70 % ethanol until they are fully coated then swiftly passing them through a flame of a Bunsen burner until the ethanol is dried, avoiding damaging/burning the leaves. Leaves were ground and suspended in 10 mM MgCl2. A 10-fold serial dilution was performed to final dilutions of 1:5, 1:50, and 1:500; 100 µl of each dilution was plated on LB agar (1.0% tryptone, 0.5% yeast extract, 1.0% NaCl, 1.3% agar) and incubated for 2-3 weeks to allow the growth of slow-growing bacteria. Colonies were then moved to fresh LB agar plates and purified. Single colonies were grown overnight in LB broth (1.0% tryptone, 0.5% yeast extract, 1.0% NaCl) for DNA extraction. DNA was extracted using the MasterPure™ Complete DNA & RNA Purification Kit (LGC Bioresearch technologies, distributed by Biozym, Hessisch Oldendorf, Germany). PCR of the isolates was done using 2 primer pairs. The primer pair 799F (5'-AACMGGATTAGATACCCKG-3') and 1193R (5'-ACGTCATCCCCACCTTCC-3') for the amplification of the V5-V7 region of the 16S rRNA region was used. The primer pair 27F (5'-AGAGTTTGATCMTGGCTCAG-3') and 1492R (5'-TACGGYTACCTTGTTACGACTT-3') amplifying V1-V9 region of the 16S rRNA gene was also used for species validation when the sequencing of the V5-V7 region deemed insufficient. PCR components were the following: 1x Q5 buffer and 0.02 U/µl Q5 polymerase (New England Biolabs, Frankfurt a.M., Germany), 200 µM forward primer, 200 µM reverse primer, and 50 ng genomic bacterial DNA. The cycling conditions were as follows: initial denaturation at 98 °C for 1 minute, 35 cycles of denaturation at 98°C for 10 seconds, annealing at 58 °C for the primer pair 799F/1193R and 63 °C for the primer pair 27F/1492R for a duration of 30 seconds, extension at 72°C for 25 seconds for the pair 799F/1193R and 60 seconds for the pair 27F/1492R. Final extension was done at 72 °C for 2 minutes. The PCR products were purified using the PCR and gel clean-up kit (Macherey Nagel, Düren, Germany) and were sent for Sanger sequencing at the Eurofins sequencing facility in Ebersberg, Germany. Bacterial isolates are listed in **Dataset S3.**

##### Amplicon sequencing of the maize phyllosphere

For amplicon sequencing i.e., the culture-independent approach and analysis of the *Z. mays* microbiome, growth conditions are as above-mentioned. Samples were collected at 15 dpi, marked by advanced pathogenesis of *U. maydis* in the inoculated maize seedlings and full maturation of tumors. The region of infection in *U. maydis* infected leaf and the corresponding area in non-infected leaves were sampled. Processing of the samples took place directly after sampling. For the sampling of whole microbiota, an average of 1.4 g of material was collected. Leaf sections were washed with sterile Milli-Q water to remove airborne contaminants and loosely attached particles, then ground with liquid Nitrogen.

As for the epiphytic fraction, we established modifications to the isolation protocol described by Li et al. (2022)[1] to account for the susceptibility of maize seedling leaves to shearing forces. These modifications allowed the collection of epiphytes by aiming to avoid disruptions of the leaf epidermis, and avoiding as much as possible the contamination of the epiphytic fraction with microbial cells spilling from the endosphere. Additionally, due to the lower DNA yield of this fraction compared to the endophyte and whole microbiota fractions (which included plant DNA), an average of 10 g of leaves were collected to meet the minimum concentration required for amplicon sequencing. Leaf sections were gently washed with sterile Milli-Q water and divided into three 50 ml Falcon tubes to facilitate and increase the efficiency of epiphyte release from the leaf surface. Samples were submerged in 40 ml of PBS buffer (pH = 7.2) supplemented with 0.01 % Tween20, and sonicated for 90 seconds at 45 kHz and 12 W in an ice bath to dislodge the epiphytes from the leaf surface. Falcon tubes containing the samples were then incubated on a rotary shaker at 150 rpm and 28 °C for 30 minutes to release the epiphytes into the PBS buffer. After incubation, leaf sections were removed from the Falcon tubes and the buffer was filtered through 0.2 µm cellulose nitrate membranes (25 mm diameter) (Whatman™, Cytiva; Merck, Darmstadt, Germany), collecting the epiphytes. The membranes containing the epiphytes were then flash frozen in liquid nitrogen.

The endophytic fraction was isolated in 2 different methods in order to ensure full and consistent representation of the endophytes. The first approach entailed the use of leaf sections that were used to isolate the epiphytes i.e., the resulting isolated microbiota here would consist of the entire endophytic fraction, plus the epiphytes that remained tightly attached to the leaves after the epiphyte isolation. The second method of collecting the endophytes was done as described in the "Culture collection of the maize phyllosphere" section**.** The sterilized leaves were then washed with sterile water to remove any DNA remnants from the dead microbial cells. A leaf imprint test was done following surface sterilization to validate the sterilization method. The leaf material from both methods was ground and flash frozen in liquid nitrogen for downstream applications.

The DNA extraction from ground leaves was done using a lysis buffer containing 100 mM Tris-HCl, 50 mM EDTA, 500 mM NaCl, 1.5 % (w/v) SDS, and 0.3 % β-mercaptoethanol followed by purification with 25:24:1 Phenol/Chloroform/Isoamyl alcohol (Carl Roth GmbH + Co. KG, Karlsruhe, Germany). Subsequent RNase digestion and protein precipitation of the DNA was done using the MasterPure™ Complete DNA & RNA Purification Kit (LGC Bioresearch technologies, distributed by Biozym, Hessisch Oldendorf, Germany).

As for the epiphytic fraction collected in the cellulose nitrate membrane, DNA isolation was performed following a custom protocol combining chemical lysis by an SDS based buffer and mechanical lysis by bead beating, followed by a phenol/chloroform/isoamyl alcohol purification protocol (Agler et al., 2016)[2].The protocol was kindly provided to us by the Kemen Lab at the University of Tübingen. In order to increase nucleic acid recovery, DNA was precipitated in 1/10 volume 3 M sodium acetate, 0.1 µg/µl molecular biology grade glycogen (Thermo Fisher Scientific, Darmstadt, Germany), and 1 volume isopropanol for 16 h at – 20 °C.

DNA quality was determined via Nanodrop 2000c spectrophotometer and quantified using Qubit 4 fluorometer (Thermo Fisher Scientific, Darmstadt, Germany). DNA samples were sent to Novogene facilities in Munich (Germany) where library preparation, sequencing, and analysis were performed according to the facility's standard method. In short, the amplification of the hypervariable region spanning V5 to V7 of 16S rRNA gene was done using the barcoded primer pair 5'- AACMGGATTAGATACCCKG-3' and 5'- ACGTCATCCCCACCTTCC-3', and the High-Fidelity Phusion polymerase. PCR products were selected, end-repaired, A-tailed, and ligated with illumina adapters. Generated libraries were pooled and sequenced on paired-end NovaSeq 6000 Illumina platform with 100K sequencing depth. Software used for analysis are summarized in Supplementary Table S3. Species annotation was performed with reference to the SILVA database [3](https://www.arb-silva.de/).

#### Infection assays

The *U. maydis* transformation assay was performed by using protoplasts prepared according to Kämper (2004)[4]. Disease assays and co-inoculations for SG200, SG200Δ*ribo1*, SG200Δ*gh25* and SG200Δ*gh25*Δ*ribo1* strains were performed according to Ökmen *et al*.[5]. In short, all *U. maydis* strains were grown in $YEPS_{Light}$ liquid medium at 28°C and 200 rpm shaking overnight. The next morning, cultures were diluted to an $OD_{600}$ of 0.25 and grown until reaching an $OD_{600}$ of 0.8–1.2. Subsequently, *U. maydis* cells were centrifuged at *3500 rpm* for 15min at RT and resuspended in 20 ml distilled water, followed by another centrifugation at *3500 rpm* for 10min. Then, pellets were resuspended in distilled water to an $OD_{600}$ of 1.0. Each *U. maydis* cell suspension was injected into stems of 6 to 7-d-old maize seedlings (GB) with a syringe with needle. Only plants were infected which had a third leaf and had no visible fourth leaf. All infection assays were performed at least in three biological replicates. In bacterial co-inoculation experiments, a final $OD_{600}$of 1.0 for *U. maydis* and 0.2 for each bacterial strain was used. To apply statistical analysis on *U. maydis* disease assays, the disease index was calculated as follows: Each plant was scored based on visible symptoms. The number of scored plants sorted into categories 'no symptoms', 'chlorosis', 'small tumor', 'normal tumor', 'heavy tumor' and 'death'. Then 'small tumor', 'normal tumor' and 'heavy tumor' were multiplied by 3, 4 and 5, respectively. All calculated numbers for each strain were summed and then divided by the total number of infected plants to obtain the disease index.

#### Simulating community growth during metabolic shifts

The complete growth medium was based on the root exudate composition reported by Horst et al. (2010)[6], with uptake bounds adapted from Schaefer et al. (2023)[7] and further adjusted to reflect tumor-associated conditions. The MICOM function complete_community_medium was applied independently to the HCom and DCom models, enforcing a community growth rate of 1 for both community models. The resulting media were merged by retaining the maximum uptake bound for overlapping metabolites. Flux balance analysis (FBA) was then run on the merged medium to confirm that both community models still achieved equal growth.

While lower growth rates could be explored directly, achieving higher growth rates required not only increasing the influx of metabolites of interest, but also removing additional growth-limiting constraints in the medium. To identify these growth-constraining metabolites and enable community model growth rates above 1, we systematically predicted growth rates for reduced uptake bound of each metabolite in the medium to 10% of its original value for both community models. Metabolites causing a growth reduction greater than 5% were classified as growth restricting. These growth restricting metabolites were partitioned into three sets: metabolites restricting both models (equal_growth_constraining_metabolites), metabolites restricting only one model (hcom or dcom_growth_constraining_metabolites), and the union of all restricting metabolites (all_constraining_metabolites). Uptake bounds for all metabolites in all_constraining_metabolites were then set to 1000 to approximate non-limiting conditions (**Fig. S10E**)

For subsequent simulations, the upper bounds of the equal_growth_constraining_metabolites were set to the highest multiplication factor used in all simulations, to remove their constraint on growth and allow growth rates above 1.

Shifts toward health- or tumor-associated conditions were simulated by reciprocally scaling the fluxes of the respective metabolite groups. For health-directed shifts, uptake bounds of tumor-associated metabolites (Gln, Asn, Arg, Ptrc, Glc, Fru) were decreased stepwise (multiplicative factors 0.9, 0.8, 0.7, etc.), while uptake bounds of health-associated metabolites were increased correspondingly (1.1, 1.2, 1.3, etc.); the inverse procedure was applied for tumor-directed shifts. At each step, FBA was performed and community growth rates were recorded and plotted against the applied scaling factors (**Fig 5).**

To test the contribution of individual groups, this procedure was repeated for individual metabolite subsets ([Gln, Asn, Arg], [Ptrc], [Glc, Fru], [$NO_3$, $NH_4$]) by varying their uptake bounds from 0 to 3 in increments of 0.1, without reciprocal scaling of other metabolite groups (**Fig S10A-D**).

### SI Tables

**Supplementary Table S1:** Taxonomic assignment of the bacterial strains of the HCom, DCom, and PdEs communities as identified by GDTB-Tk[8–14] (ANI based analysis) and TYGS (dDDH d4-based analysis)[15–22]

| Community | GTDB-Tk taxonomy | ANI to closest GTDB genome (%) | Closest GTDB reference | TYGS taxonomy | dDDH to closest type strain (%) | Final assignment |
|---|---|---|---|---|---|---|
| **HCom** | *Priestia megaterium* | 97.32 | GCF_000832985.1 | *Priestia megaterium* | 77.7 | *Priestia megaterium* |
| **HCom** | *Staphylococcus epidermidis* | 99.62 | GCF_006742205.1 | *Staphylococcus epidermidis* | 96.6 | *Staphylococcus epidermidis* |
| **HCom** | *Arthrobacter sp.* | N/A | N/A | *Arthrobacter sp. [potential new species]* | N/A | *Arthrobacter sp.* ***[potential new species]]*** |
| **HCom** | *Paenibacillus amylolyticus* | 98.6 | GCF_008386395.1 | *Paenibacillus sp. [potential new species]* | 57.7 | *Paenibacillus amylolyticus* |
| **HCom** | *Bacillus licheniformis* | 99.72 | GCF_000011645.1 | *Bacillus licheniformis* | 97.9 | *Bacillus licheniformis* |
| **HCom** | *Peribacillus sp030348945* | 98.66 | GCF_050279365.1 | *Peribacillus sp.* ***[potential new species]*** | N/A | *Peribacillus sp.* |
| **DCom** | *Enterobacter kobei* | 99.32 | GCF_001729765.1 | *Enterobacter kobei* | 94.7 | *Enterobacter kobei* |
| **DCom** | *Brevibacterium sediminis* | 97.53 | GCF_013623905.1 | *Brevibacterium sediminis* | 77.6 | *Brevibacterium sediminis* |
| **DCom** | *Microbacterium sp.* | N/A | N/A | *Microbacterium sp.* ***[potential new species]*** | N/A | *Microbacterium sp.* ***[potential new species]*** |
| **DCom** | *Erwinia aphidicola* | 98.15 | GCF_024169515.1 | *Erwinia aphidicola* | 97.2 | *Erwinia aphidicola* |
| **DCom** | *Klebsiella ornithinolytica* | 99.59 | GCF_001598295.1 | *Klebsiella ornithinolytica* | 97.2 | *Raoultella ornithinolytica* (***reclassification of Klebsiella ornithinolytica***) |
| **DCom** | *Chryseomonas aestiva* | 96.86 | GCF_039908605.1 | *Pseudomonas aestiva* | 72.6 | *Pseudomonas aestiva* (***reclassification of Chryseomonas aestiva)*** |
| **PdEs** | *Pantoea dispersa* | 98.2 | GCF_014155765.1 | *Pantoea dispersa* | 84.1 | *Pantoea dispersa* |
| **PdEs** | *Enterobacter sichuanensis* | 98.13 | GCF_002939185.1 | *Enterobacter sichuanensis* | 84.7 | *Enterobacter sichuanensis* |

**Supplementary Table S2:** Summary of metabolite shifts associated with *U. maydis* tumor formation in maize leaves, summarized from Horst et al. (2010)[6]. Arrows indicate the direction of change reported for tumor tissue relative to healthy leaf tissue in Horst et al. (2010), based on qualitative assessment of metabolite abundances across figures and text. Directionality reflects consistent trends across metabolites within a class and does not imply quantitative fold changes.

| Metabolite class | Representative metabolites | Tumor vs healthy | Basis in Horst et al. (2010) |
|---|---|---|---|
| **Reduced organic nitrogen** | Glutamine, asparagine, arginine | ↑ | Increased pools reported in metabolite profiling |
| **Total free amino acids** | Multiple | ↑ | Consistent elevation across amino acid classes |
| **Polyamines** | Putrescine | ↑ | Reported accumulation in tumor tissue |
| **Inorganic nitrate** | $NO_3^-$ | ↓ | Reduced nitrate content in tumors |
| **Inorganic ammonium** | $NH_4^+$ | ↓ | Lower free ammonium pools |
| **Soluble sugars** | Glucose, fructose | ↑ | Increased sugar content in tumors |

**Supplementary Table S3:** Summary of software and tools used for the analysis of the amplicon sequencing data.

| Software and tools used | Identifier | Source |
|---|---|---|
| **QIIME2 (202202)** | https://qiime2.org/ | Bolyen et al., 2019[23] |
| **QIIME2 DADA2** | http://benjjneb.github.io/dada2/ | Callahan et al., 2016[24] |
| **FLASH (1.22.11)** | http://ccb.jhu.edu/software/FLASH | Magoč et al., 2011[25] |
| **fastp (0.23.1)** | https://github.com/OpenGene/fastp | Chen et al., 2025[26] |
| **vsearch (2.16.0)** | http://github.com/torognes/vsearch | Edgar et al., 2011[27] |
| **Perl (5.26.2) with SVG function** | https://perl.org, https://metacpan.org/pod/SVG | |
| **R (4.0.3) v with pheatmap() function** | https://www.r-project.org/, https://cran.r-project.org/web/packages/pheatmap/index.html | |
| **QIIME2 (202202) for alpha diversity (dominance, Shannon)** | https://scikit.bio/docs/dev/generated/skbio.diversity. alpha.dominance.html#r321a44067455-1 https://scikit.bio/docs/dev/generated/skbio.diversity. alpha.shannon.html | Bolyen et al., 2019[23] |
| **R V.0.3 with Tax4Fun (V0.3.1) package** | https://www.r-project.org/, https://tax4fun.gobics.de/ | Aßhauer et al., 2015[28] |
| **Python (3.6.13) with FAPROTAX tool** | https://www.python.org/, http://www.loucalab.com/archive/FAPROTAX | Louca et al., 2016[29] |

#### SI References


1. Li, P. D. *et al.* The phyllosphere microbiome shifts toward combating melanose pathogen. *Microbiome* **10**, 56- (2022).
2. Ruhe, J. *et al.* Obligate biotroph pathogens of the genus albugo are better adapted to active host defense compared to niche competitors. *Front. Plant Sci.* **7**, 184119 (2016).
3. Quast, C. *et al.* The SILVA ribosomal RNA gene database project: improved data processing and web-based tools. *Nucleic Acids Res.* **41**, D590–D596 (2013).
4. Kämper, J. A PCR-based system for highly efficient generation of gene replacement mutants in Ustilago maydis. *Mol. Genet. Genomics* (2004) doi:10.1007/s00438-003-0962-8.
5. Ökmen, B. *et al.* A conserved extracellular Ribo1 with broad-spectrum cytotoxic activity enables smut fungi to compete with host-associated bacteria. *New Phytol.* **240**, 1976–1989 (2023).
6. Horst, R. J. *et al.* Ustilago maydis infection strongly alters organic nitrogen allocation in maize and stimulates productivity of systemic source leaves. *Plant Physiol.* **152**, 293–308 (2010).
7. Schäfer, M. *et al.* Metabolic interaction models recapitulate leaf microbiota ecology. *Science (80-. ).* **381**, 42–55 (2023).
8. Chaumeil, P. A., Mussig, A. J., Hugenholtz, P. & Parks, D. H. GTDB-Tk v2: memory friendly classification with the genome taxonomy database. *Bioinformatics* **38**, 5315–5316 (2022).
9. Parks, D. H. *et al.* GTDB: an ongoing census of bacterial and archaeal diversity through a phylogenetically consistent, rank normalized and complete genome-based taxonomy. *Nucleic Acids Res.* **50**, D785–D794 (2022).
10. Matsen, F. A., Kodner, R. B. & Armbrust, E. V. pplacer: linear time maximum-likelihood and Bayesian phylogenetic placement of sequences onto a fixed reference tree. *BMC Bioinformatics* **11**, 538 (2010).
11. Shaw, J. & Yu, Y. W. Fast and robust metagenomic sequence comparison through sparse chaining with skani. *Nat. Methods 2023 2011* **20**, 1661–1665 (2023).
12. Hyatt, D. *et al.* Prodigal: Prokaryotic gene recognition and translation initiation site identification. *BMC Bioinformatics* **11**, 119- (2010).
13. Price, M. N., Dehal, P. S. & Arkin, A. P. FastTree 2 – Approximately Maximum-Likelihood Trees for Large Alignments. *PLoS One* **5**, e9490 (2010).
14. Eddy, S. R. Accelerated Profile HMM Searches. *PLOS Comput. Biol.* **7**, e1002195 (2011).
15. Meier-Kolthoff, J. P. & Göker, M. TYGS is an automated high-throughput platform for state-of-the-art genome-based taxonomy. *Nat. Commun. 2019 101* **10**, 2182- (2019).
16. Meier-Kolthoff, J. P., Carbasse, J. S., Peinado-Olarte, R. L. & Göker, M. TYGS and LPSN: a database tandem for fast and reliable genome-based classification and nomenclature of prokaryotes. *Nucleic Acids Res.* **50**, D801–D807 (2022).
17. Freese, H. M., Meier-Kolthoff, J. P., Sardà Carbasse, J., Afolayan, A. O. & Göker, M. TYGS and LPSN in 2025: a Global Core Biodata Resource for genome-based classification and nomenclature of prokaryotes within DSMZ Digital Diversity. *Nucleic Acids Res.* **54**, D884–D891 (2026).
18. Meier-Kolthoff, J. P., Auch, A. F., Klenk, H. P. & Göker, M. Genome sequence-based species delimitation with confidence intervals and improved distance functions. *BMC Bioinformatics* **14**, (2013).
19. Meier-Kolthoff, J. P. *et al.* Complete genome sequence of DSM 30083T, the type strain (U5/41T) of Escherichia coli, and a proposal for delineating subspecies in microbial taxonomy. *Stand. Genomic Sci.* **9**, 2- (2014).
20. Lefort, V., Desper, R. & Gascuel, O. FastME 2.0: A Comprehensive, Accurate, and Fast Distance-

Based Phylogeny Inference Program. *Mol. Biol. Evol.* **32**, 2798–2800 (2015).

21. Farris, J. S. Estimating Phylogenetic Trees from Distance Matrices. *https://doi.org/10.1086/282802* **106**, 645–668 (1972).

22. Kreft, L., Botzki, A., Coppens, F., Vandepoele, K. & Van Bel, M. PhyD3: a phylogenetic tree viewer with extended phyloXML support for functional genomics data visualization. *Bioinformatics* **33**, 2946–2947 (2017).

23. Bolyen, E. *et al.* Reproducible, interactive, scalable and extensible microbiome data science using QIIME 2. *Nat. Biotechnol. 2019 378* **37**, 852–857 (2019).

24. Callahan, B. J. *et al.* DADA2: High-resolution sample inference from Illumina amplicon data. *Nat. Methods 2016 137* **13**, 581–583 (2016).

25. Magoč, T. & Salzberg, S. L. FLASH: fast length adjustment of short reads to improve genome assemblies. *Bioinformatics* **27**, 2957–2963 (2011).

26. Chen, S. fastp 1.0: An ultra-fast all-round tool for FASTQ data quality control and preprocessing. *iMeta* **4**, e70078 (2025).

27. Rognes, T., Flouri, T., Nichols, B., Quince, C. & Mahé, F. VSEARCH: A versatile open source tool for metagenomics. *PeerJ* **2016**, e2584 (2016).

28. Aßhauer, K. P., Wemheuer, B., Daniel, R. & Meinicke, P. Tax4Fun: predicting functional profiles from metagenomic 16S rRNA data. *Bioinformatics* **31**, 2882 (2015).

29. Louca, S., Parfrey, L. W. & Doebeli, M. Decoupling function and taxonomy in the global ocean microbiome. *Science (80-. ).* **353**, 1272–1277 (2016).
